# Epigenome and interactome profiling uncovers principles of distal regulation in the barley genome

**DOI:** 10.1101/2025.02.27.640517

**Authors:** Pavla Navratilova, Simon Pavlu, Zihao Zhu, Zuzana Tulpova, Ondrej Kopecky, Petr Novak, Nils Stein, Hana Simkova

**Affiliations:** Institute of Experimental Botany of the Czech Academy of Sciences, Olomouc, Czech Republic; Department of Cell Biology and Genetics, Faculty of Science, Palacky University, Olomouc, Czech Republic; Leibniz Institute of Plant Genetics and Crop Plant Research Gatersleben, Germany; Czech Academy of Sciences, Institute of Plant Molecular Biology, Ceske Budejovice, Czech Republic; Crop Plant Genetics, Martin Luther University of Halle-Wittenberg, Halle (Saale), Germany

**Keywords:** epigenetics, interactome, chromatin states, *Hordeum vulgare*, Morex, embryo, regulome, *Vrn3*, LEA genes, epigenome browser

## Abstract

Regulation of transcription initiation is the ground level of modulating gene expression during plant development. This process relies on interactions between transcription factors and *cis-* regulatory elements (CREs), which become promising targets for crop bioengineering. To annotate CREs in the barley genome and understand mechanisms of distal regulation, we profiled several epigenetic features across three stages of barley embryo and leaves, and performed HiChIP to identify activating and repressive genomic interactions. Using machine learning, we integrated the data into seven chromatin states, predicting ∼77,000 CRE candidates, collectively representing 1.43% of the barley genome. Identified genomic interactions, often spanning multiple genes, linked thousands of CREs with their targets and revealed notably frequent promoter-promoter contacts. Using the LEA gene family as an example, we discuss possible roles of these interactions in transcription regulation. On the Vrn3 gene, we demonstrate the potential of our datasets to predict CREs for other developmental stages.

## Introduction

Eukaryotic gene expression is regulated at multiple levels, with transcription initiation being a foundational level governed by a *cis*-regulome. This includes core promoters, binding RNA polymerase pre-initiation complex, and proximal and distal elements^1^. The distal *cis*-regulatory elements (CREs) can function as enhancers, silencers, or insulators. The ratio of proximal to distal CREs varies depending on the intergenic space, with larger genomes incorporating more distal, chromatin loop-mediated regulation^2^.

In this context, barley (*Hordeum vulgare* L.), one of the earliest domesticated crops, represents a good experimental model for small-grain temperate-zone cereals due to its diploid genome of almost 5 Gb^3,4^. Barley core promoters were analyzed in detail in our previous work^5^, but knowledge about the localization and function of proximal and distal CREs remains scarce. The barley genome has vast intergenic spaces, which are likely to be rich in the distal CREs, as indicated by the distribution of epigenetic features associated with transcriptional activity^6^. Comprehensive studies including whole-genome profiling of various epigenetic features have been published for several cereal species, including bread wheat^7^, maize^8^, and rice^10^. Such analyses, typically conducted using seedlings or leaves, provided insights into the general regulatory potential of CREs in terminally differentiated tissues. Given that *cis-*regulation is most intense during embryonic development^11^, current CRE collections are unlikely to be complete and need to be extended to include those from actively differentiating stages.

In contrast to promoters and proximal elements, located just upstream of transcription start sites (TSSs), distal CREs, lacking recognizable sequence signatures and residing at up to 1 Mb from their target^12^, are notoriously difficult to locate. Since their activity is closely linked to epigenetic features such as chromatin accessibility, histone modifications, and DNA methylation, the combination of these marks serves as a guide to their annotation. DNA methylation is a stable CRE characteristic in plants with many elements remaining unmethylated even outside of their activity time window^13^. In contrast, accessible chromatin regions (ACRs), identified by ATAC-seq, are more dynamic, reflecting immediate transcription factor (TF) binding, followed by histone H3/H4 acetylation, RNA polymerase association and H3K4 methylation, as reviewed in^14,15^. Opposed to that, H3K27me3, a result of Polycomb activity, decorates histones in facultatively repressed regions, as reviewed in^16^. Collectively, specific combinations of histone modifications, known as the histone code^17^, with ACRs and unmethylated regions are determinants of the current functional status of a given chromatin segment, efficiently integrated by computational methods based on machine learning^18,19^.

Only a handful of studies have comprehensively defined the CREs including their targets^8,20–23^. When a CRE is distal to its target promoter, it establishes a transient physical interaction with the target gene via chromatin looping^14,24–26^. In Polycomb gene silencing, H3K27me3-associated loops between genes and their enhancers, or ’silencing hubs‘, gather genes and CREs to repress their activity during development^27^. Chromatin Conformation Capture (3C)-based techniques remain the only experimental method to determine CRE target genes^28^. Their efficiency improves when combined with antibody subtraction (HiChIP) or promoter capture (Capture Hi-C)^29,30^, as reviewed in^31^.

Chromatin loop mediation is one of the roles of unstable small non-coding RNAs transcribed from animal enhancers (enhancer RNA; eRNA)^32^. Tens of thousands of these RNAs, transcribed uni- or bidirectionally, have been detected in mammalian cells, where they also facilitate chromatin remodeling and transcription factor recruitment^32,33^. However, the existence and function of eRNAs in plants remains debated^34^. Plant CRE-associated RNAs are predominantly unidirectional, with bidirectional transcripts much less common compared to vertebrates^35^. Previously, we detected stable capped transcripts genome-wide in barley embryos by CAGE^5^, but the sensitivity for low-expressed, unstable RNA was insufficient, calling for further study.

An increasing number of studies provide evidence of CREs overlapping with eQTL and their association with agronomically important traits in crops^22,36^. Understanding transcription *cis-* regulation during embryonic stages expands our knowledge of the establishment of valuable traits such as seed vigor and seed longevity^37^. LATE EMBRYOGENESIS ABUNDANT (LEA) proteins play a key role in these processes, as reviewed by^37^. Expressed in response to water loss during seed maturation as well as during vegetative growth, they contribute to desiccation tolerance and adaptation to drought stress^38^. LEA genes are evolutionarily conserved among angiosperms, with the *LEA_5* family showing the highest conserved synteny, indicating evolutionary constraints on maintaining the integrity of their genomic context^38^. This, along with the clustered distribution of *LEA_5* genes observed across the barley pangenome v2 collection^39^, highlights them as promising targets for investigating their interactome and epigenomic context.

Another fundamental process in plant development is the transition from the vegetative to the reproductive stage, regulated by vernalization-related genes and miRNAs in response to environmental cues. The vernalization involves epigenetic mechanisms, including changes of histone modifications and 3D chromatin remodelling in winter wheat^40,42^. One of the crucial regulators, the product of *Vernalization 3* (*Vrn3*), putatively orthologous to Arabidopsis *FLOWERING LOCUS T* (*FT*), functions as florigen in wheat^43^. Its expression is controlled not only by a vernalization- but also by an age-related pathway, ensuring that flowering occurs in adulthood^40,41^. Two transcriptional enhancers were found upstream of *Vrn3* in winter wheat^40^, raising a question of whether the same CREs could control *Vrn3* transcription in spring barley.

Here, we predict the *cis-*regulatory landscape of the barley genome in the developing embryo, germinating embryo and leaf. For each stage, we generated and integrated whole-genome profiles of several epigenetic features and complemented them by interactome data for the maturing embryo. Our study concluded with a comprehensive map of key epigenome features, predicted CRE candidates (cCREs), and genomic interactions. Analysis of these interactions assigned gene targets to multiple identified cCREs and revealed diverse interaction classes. Nascent RNA sequencing confirmed the minor role of non-coding transcription in CRE activity. On a cluster of LEA genes, we demonstrated interactions between promoters and discussed their role in transcriptional regulation. At the *Vrn3* locus, we illustrate the power of our datasets to predict distal CREs for other developmental stages. This integrative analysis advances the understanding of transcription regulation in large plant genomes and provides a valuable resource for bioengineering in barley. All data visualizations will be made available through an interactive genome browser (https://olomouc.ueb.cas.cz/en/resources/barleyepibase).

## Results

### Profiles of epigenetic features enable annotation of *cis-*regulatory elements

To enable comprehensive annotation of the barley *cis-*regulome, we followed a workflow outlined in Fig. 1A. We selected a minimal set of epigenetic features and generated genomic profile datasets from three stages of barley embryo development – eight days after pollination (8DAP), 24 days after pollination (24DAP), and four days of germination (4DAG) – as well as from young leaf tissue. High quality of our datasets was proven by standard profile distributions around annotated gene TSSs (Fig. 1B) and documented through peak counts and replicate overlaps, summarized in Suppl. Table 1. The active TSSs are enriched with histone modification marks and ATAC-seq signals, while being devoid of DNA methylation, as expected^14^.

**Fig. 1.**
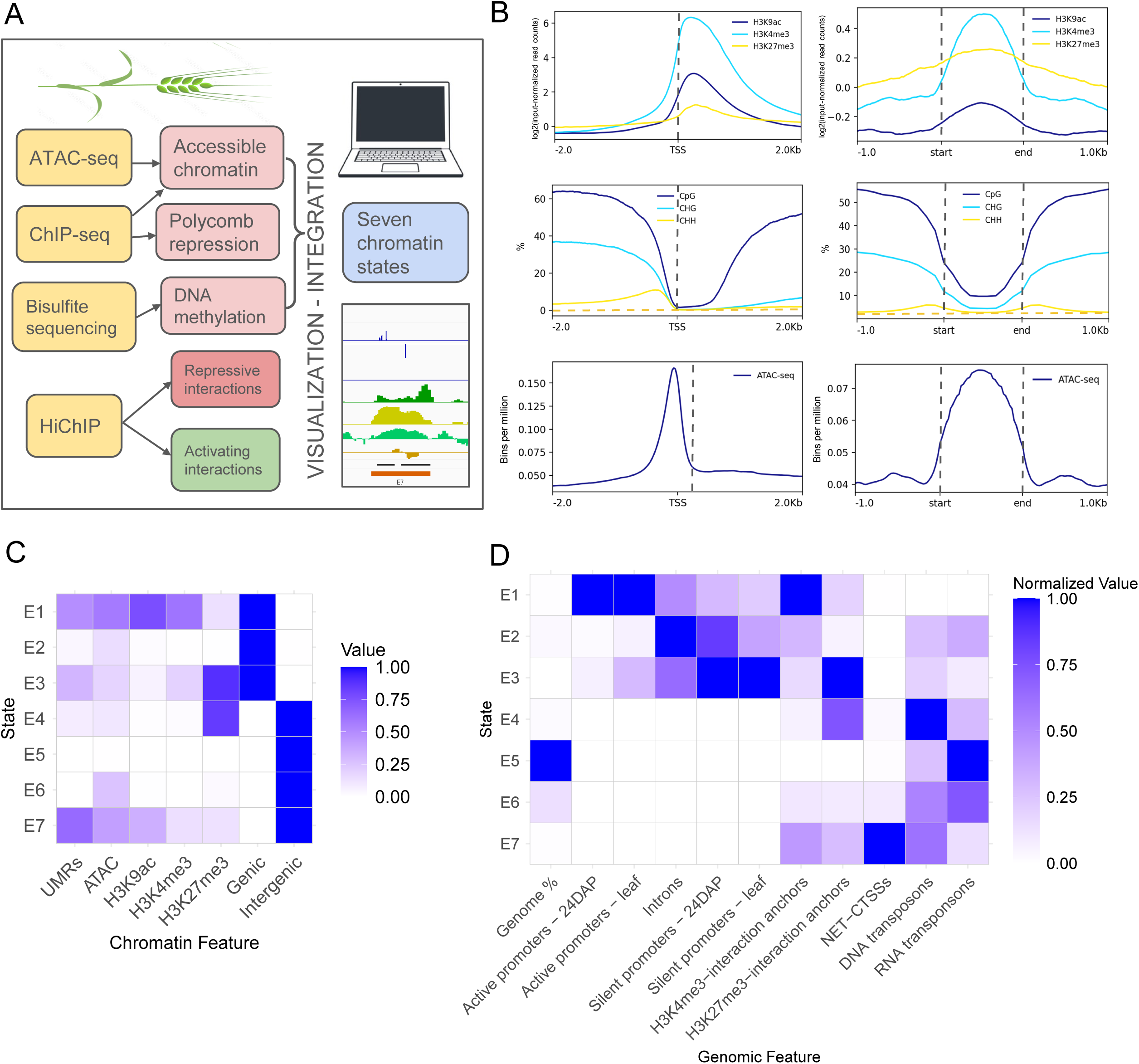
Key epigenetic chromatin features facilitate annotation of *cis*-regulatory elements. (A) Components of the barley *cis*-regulome and interactome analysis. (B) Profiles of coverages of key epigenetic features – key histone modifications (top), DNA methylation (middle), and open chromatin (bottom) around active TSSs (left) and across segments with chromatin state 7 in the 24DAP embryo (right). (C) A Chromatin State Emission Values heatmap for the ChromHMM model distinguishing seven chromatin states. The intensity of the blue color corresponds to emission values, indicating the likelihood of a given state being associated with the chromatin feature. (D) Overlap enrichment heatmap shows the fold enrichment of each state of the segmentation from (C) for a set of selected 24DAP genomic features. H3K4me3 and H3K27me3 interaction anchors correspond to 5-kb interacting bins from HiChIP analysis, while NET-CTSS corresponds to clusters of nascent capped transcript initiation used for eRNA detection.

A notable CRE feature that appears largely independent of cellular context is DNA methylation. We quantified the methylation levels in 24DAP embryos (Suppl. Fig. 1A, B) and leaf tissue (BS-seq data from^44^. Consistent with findings in other cereals, barley exhibits a highly methylated genome, with average genome-wide methylation levels of 88.6%, 58.1%, and 1.4% in the CpG, CHG, and CHH sequence contexts, respectively. We defined unmethylated regions (UMRs) in the 24DAP and leaf samples and subtracted all UMRs overlapping with genic regions, defined as described below, resulting in 102,287 and 102,362 UMRs, respectively. We then identified ‘permanent’ UMRs as the overlap between these two sets, yielding a total of 74,614 intergenic UMRs.

As a cell-specific feature that serves as a useful proxy for functional sequences associated with transcriptional activity, we assessed open chromatin by ATAC-seq across all four stages (Suppl. Fig. 1C). Additionally, we immunoprecipitated chromatin using antibodies against three histone posttranslational modifications to capture actively transcribed regions (H3K4me3 and H3K9ac) and Polycomb-repressed genomic regions (H3K27me3) (Suppl. Fig. 2).

Importantly, to prevent contamination of our *cis*-regulome analysis with unannotated genes (Suppl. Fig. 3) and to avoid false positives in defining CRE candidates, which resemble promoters in their epigenetic features, we carefully assessed the protein- and lncRNA-coding potential based on RNA-seq data and a previous publication^45^. Such transcribed regions, extended by 500 bp in both directions, along with the high- and low- confidence MorexV3 gene annotations, extended by 500 bp at the TSS, define the ‘genic’ portion of the genome. The complementary genomic regions were classified as ‘intergenic’, and both categories, essential for the annotation of potential regulatory elements, were subsequently used to define coding potential in the chromatin state model.

Discovering *de novo* the major recurring combinatorial and spatial chromatin patterns, known as ‘chromatin states’, allows for narrowing down putative regulatory elements. To achieve this, we integrated ATAC-seq, ChIP-seq, and UMR data from four stages of barley development with coding-potential segmentation and conducted chromatin state analysis using ChromHMM^46^. This approach binarizes the data and applies a multivariate hidden Markov model to learn a defined number of chromatin states. The resulting minimal model, which resolved both active and silent, genic and intergenic situations at a 200-bp resolution, comprised seven chromatin states, referred to as E1-7 (Fig. 1C). Following this, we conducted genomic feature overlap analysis for 24DAP (Fig 1D) and leaf (Suppl. Fig. 4A) samples to quantify normalized overlaps between individual states and various genomic features.

The resulting probability heatmaps confirmed the functional identity of genomic segments defined by the seven states and uncovered transcription-related dynamics of promoter states between stages. The active chromatin state E1, enriched in H3K4me3 and H3K9ac and characterized by open chromatin and UMRs, is predominantly associated with active gene promoters. In contrast, the repressed coding state E3 overlaps especially with Polycomb-silenced promoters and genes, marked by H3K27me3. Importantly, our chromatin analysis identified largely unmethylated intergenic regions dominated by open chromatin and acetylated histones with a certain likelihood of histone H3K4 and H3K27 tri-methylations, corresponding to state E7, as confirmed by chromatin profile distributions across E7 segments (Fig. 1B). Pool of 77,383 E7-state segments from four analyzed stages, likely enriched in active CREs, covers 1.43% of barley MorexV3 genome. Stage-specific E7 segments (Suppl. File 1) were used for further analyses. The overlap of E7 segments across samples revealed their dynamics: 63.4% of all elements were commonly detected in all stages, whereas 2.4% to 9.6% were stage-specific (Fig. 2A). The H3K27me3-enriched state E4 segments represent intergenic Polycomb-silenced regions, with a mean size of 3.5 kb, compared to the 800 bp E7 segments. Approximately 76% of E4 segments embedded relatively small UMRs, with a mean size of 400 bp, suggesting their function in CRE silencing.

**Fig. 2.**
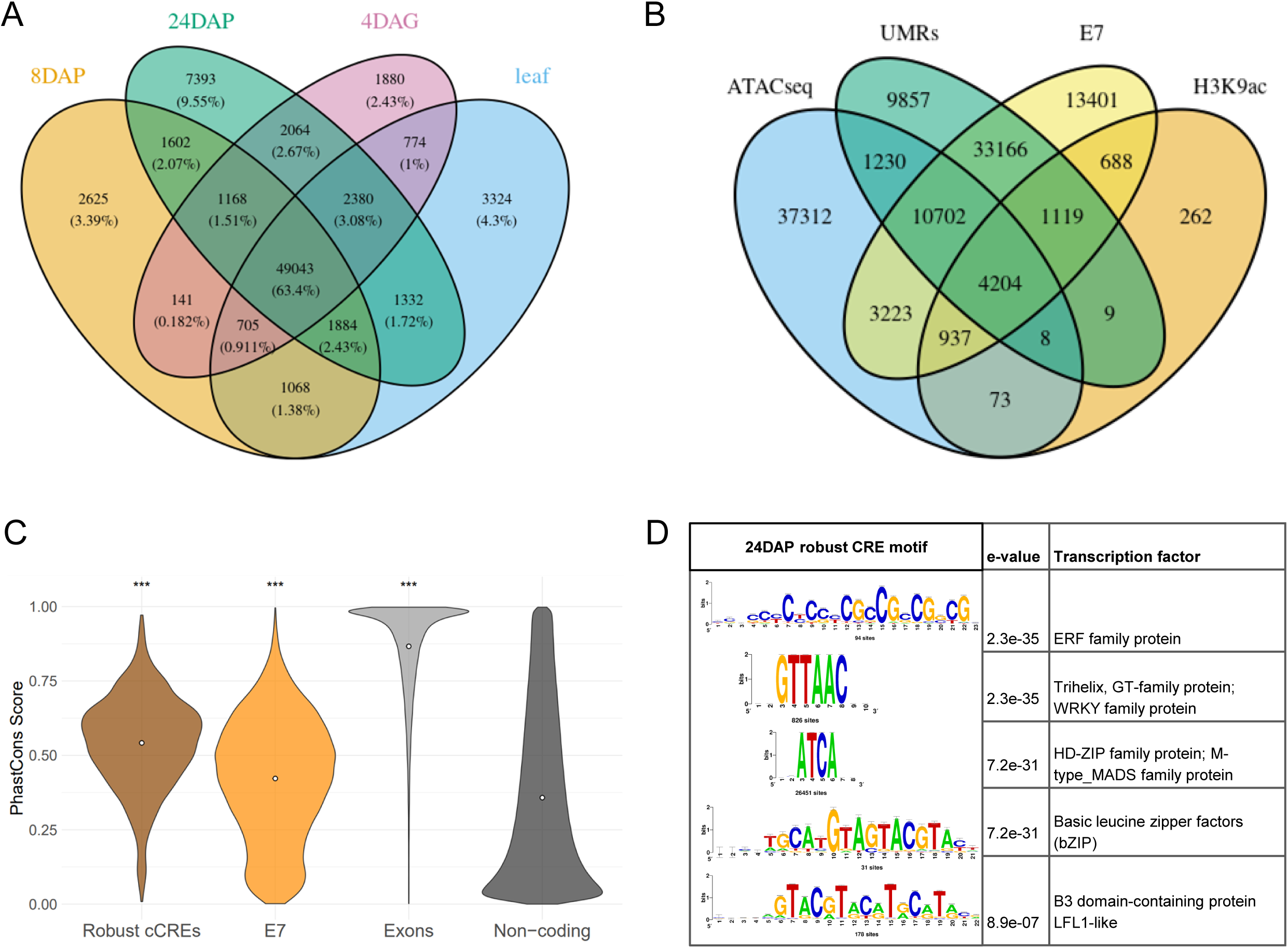
Characterization of candidate *cis*-regulatory elements. (A) Venn diagram showing the dynamics of E7 segments across four developmental stages - embryo at 8DAP, 24DAP and 4DAG, and young leaf tissue. (B) The overlap of activating genomic features (peaks of ATAC- seq and H3K9ac, and UMRs) with E7 segments, corresponding to the middle section of the Venn diagram, defines the robust cCRE set, shown here for the 24DAP embryo. (C) Distribution of sequence conservation scores across E7 segments and robust cCREs, compared to exons of high-confidence genes and non-coding regions, shows that both robust cCREs and E7 segments tend to be conserved across cereal genomes. The white dots indicate mean values for each set. Significances of distribution differences compared to the binned non-coding genome were tested using Wilcoxon test (p-values < 0.005 labelled by ***). (D) Transcription factor binding sites enriched in the 24DAP-specific robust cCREs dataset.

To narrow down a set of high-confidence active CRE candidates for each stage, we intersected stage-specific intergenic ATAC-seq peaks with UMRs and H3K9ac. The resulting stage-specific sets, each covering ∼0.1% of MorexV3 genome, are referred to as ‘robust cCREs’ and comprise 3,970-6,542 elements (Fig. 2B, Suppl. Fig. 4C). The vast majority of the robust cCREs overlap with E7 segments, while subsets of the E7 segments lacking overlap with ATAC-seq or H3K9ac peaks exhibit weaker E7 chromatin feature profiles (Suppl. Fig. 4D), highlighting the higher sensitivity of ChromHMM compared to peak calling in identifying regulatory elements.

Evolutionary conservation of non-coding sequences is one of the key indicators for identifying regulatory elements^47^. To assess conservation of our predicted regulatory elements, we calculated per-base sequence conservation from whole-genome multiple alignment of five grass species (*Hordeum vulgare*, *Triticum urartu*, *Secale cereale*, *Brachypodium distachyon*, and *Aegilops tauschii*). The distributions of average PhastCons scores of the robust cCREs and E7 elements were significantly higher than those of non-coding regions but still lower than those of exons (Wilcoxon test, all p-values < 2.2e-16), pointing to the functional importance of the predicted regulatory elements (Fig. 2C).

Another distinctive feature of CRE sequences is their enrichment in transcription factor binding sites (TFBSs). We analyzed TFBS content in robust cCREs through motif enrichment analysis using RSAT^48^, identifying sets of developmental and hormone-responsive factor binding sites. At 24DAP, the Ethylene-responsive factor (ERF) motif was the most significant, consistent with its overexpression in embryos and role in starch formation^49^. Other highly enriched motifs belonged to transcription factors crucial for embryonic development^11^, including Trihelix, GT- family, HD-ZIP, MADS-family, bZIP, and B3 domain-containing LFL-like proteins (Fig. 2D). In 8DAP-specific elements, the GAGA-motif binding transcription factor BPC-like, required for seed development via homeotic TF regulation^50^, was highly significant (Suppl. Fig. 5A). In the 4DAG sample, the most significant motif was that of ERF (Suppl. Fig. 5B), followed by ZFHD10-3- and NAC factor-binding sequences. In leaf robust cCREs, ERF was again the most enriched, followed by Wuschel-like homeobox and B3 domain-containing protein. These findings underscore the functional importance of the identified robust cCREs in barley growth and development.

### Barley bidirectional and unidirectional unstable transcripts are an infrequent feature associated with cCREs

Active CREs may produce unstable transcription signals (eRNA), both unidirectional and bidirectional. To detect 5’ capped short unstable eRNA species in the 4DAG embryo, we utilized native elongating transcript–cap analysis of gene expression (NET-CAGE)^51^. From the resulting data, summarized in Suppl. Table 2, we identified 72,119 NET-CAGE tag clusters, with their dominant tags indicating the position of TSSs, which co-localized fairly well with the annotated gene TSSs positions (Fig. 3A, C). The NET-CAGE cluster annotation (Fig. 3D) defined 16,592 intergenic clusters. Filtering against the previously generated ‘genic’ portion of the genome, left us with 8,122 strictly intergenic NET-CAGE clusters, which we analyzed for transcript directionality, finding that 5,160 were unidirectional (U), 1,669 bidirectional (B) and only 595 clusters were bidirectional and balanced - a configuration attributed to canonical eRNA in animals. The remaining bidirectional clusters appeared unbalanced (BU), with TPM values different for the signal pairs (Fig. 3B). We found that particularly the BU and U NET-CAGE clusters were highly overrepresented (randomization test) in our robust cCRE set as well as in state E7, unlike in the E4 genomic segments (Fig. 3E). Nevertheless, the NET-CAGE signal was observed only in ∼10% (460/4,396) robust cCREs and 3% (1,748/60,605) E7 segments, suggesting that transcription of eRNA is not a typical attribute of barley CREs.

**Fig. 3.**
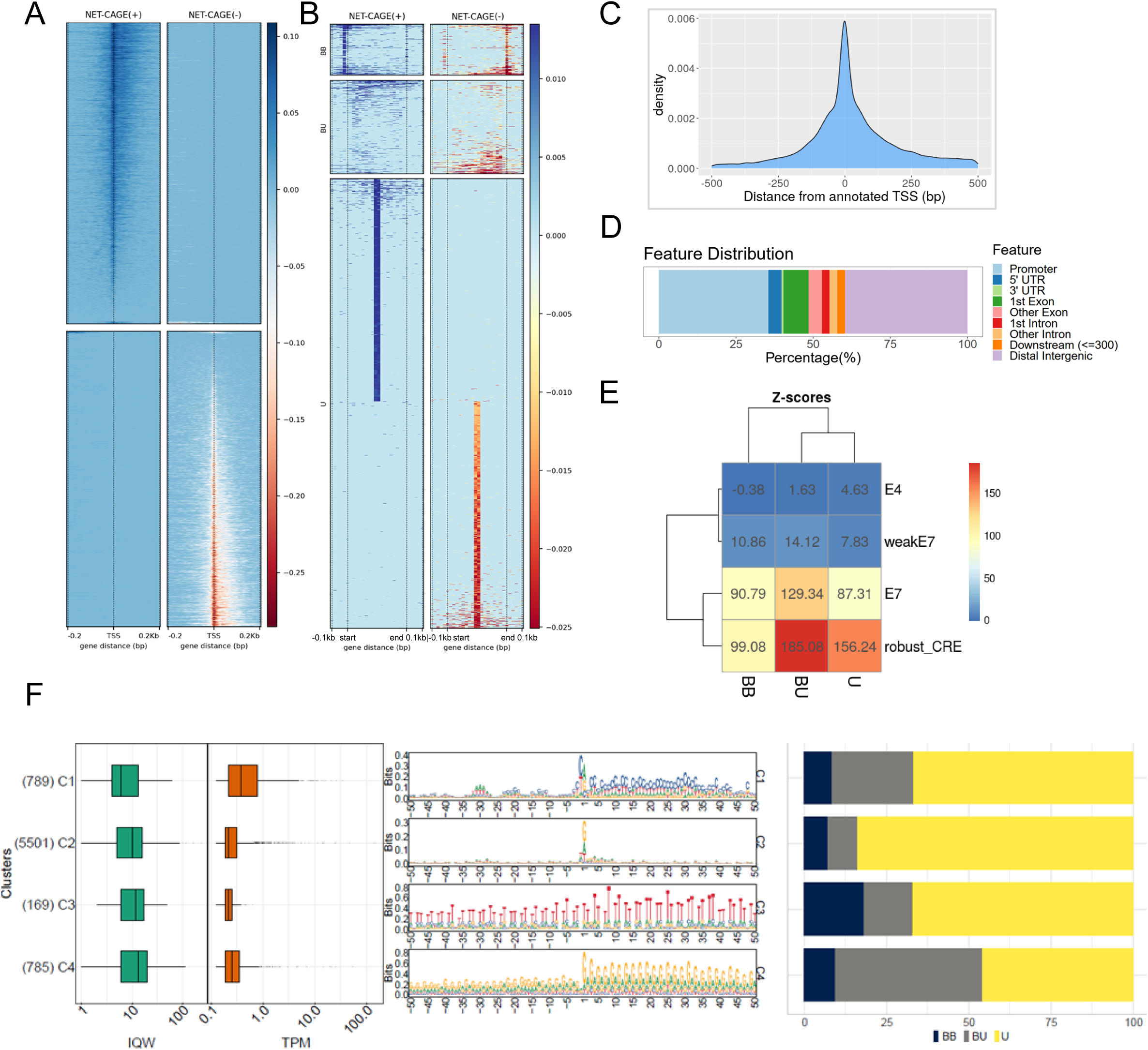
Features of native non-genic transcripts identified by NET-CAGE in the 4DAG embryo. (A) Heatmap of NET-CAGE cluster distributions on both DNA strands around annotated TSSs, indicating strong association and the absence of upstream antisense RNA signals. (B) Heatmap showing the three main directionality patterns of the non-coding RNA (ncRNA). The most abundant category consists of unidirectional transcripts resembling protein-coding gene promoters. (C) Distribution of dominant NET-CAGE tags around annotated (MorexV3) gene TSSs. (D) Genomic feature annotation of NET-CAGE clusters. (E) Overlap of the three ncRNA-directionality patterns with E4 and E7 state segments and robust cCREs. The resulting Z-scores from the randomization test of feature overlap reveal a strong overrepresentation of ncRNA in the robust cCRE and E7 sets. (F) Four main clusters of sequence architectures, together with Inter-Quartile Width (IQW) and TPM (tags per million) values. Dark blue, grey, and yellow bars represent the composition of architecture clusters based on the three directionality groups (BB = bidirectional balanced, BU = bidirectional unbalanced, U = unidirectional).

Taking advantage of the single-base precision of the dominant TSS positions, we addressed the sequences flanking the initiators of these transcripts. Using the seqArchR clustering algorithm, we identified four distinct sequence architectures at the TSS (Fig. 3F) and contextualized them with Inter-Quartile Width (IQW) and tags per million (TPM) values and the three directionality groups. The first cluster (C1) resembles coding-gene core promoters with a canonical CA initiator and a hint of a TATA-box approximately 35 bp upstream, but exhibits TPM values <1. This suggests that these may be promoters of lowly expressed unannotated genes or enhancers with a promoter-like architecture. The second, most abundant cluster (C2) resembles a promoter starting from a non-canonical G-initiator while lacking other conserved motifs. The remaining two clusters (C3 and C4) represent low-complexity, microsatellite-like sequences. These clusters show a slightly higher tendency for bidirectional transcription compared to the promoter-like clusters, which are predominantly transcribed in one direction.

### Histone modification-centric chromatin conformation capture detects activating and repressive interactions

To identify spatial chromatin interactions indicative of contacts between genes and their CREs, we performed HiChIP using anti-H3K4me3 and anti-H3K27me3 antibodies, to enrich for activating and silencing interactions, respectively. The experiments were conducted on G1- phase nuclei prepared from 24DAP embryos in two replicates, which yielded highly correlating data (Suppl. Fig. 6A). The HiChIP signal enrichment at ChIP-seq peaks indicated good sample quality (Suppl. Fig. 6B). The resulting valid interaction pool was analyzed using FitHiChIP^52^ to identify pairs of regions (‘anchors’) with a significant number of reads mapping between them, representing biologically meaningful chromatin interactions (Fig. 4A, B). The FitHiChIP ‘peak-to-all’ mode enabled the identification of cCREs lacking H3K4me3/H3K27me3 that interacted with marked promoters. For H3K4me3 at the 5-kb resolution, which is the dataset used for the majority of the analyses, the average and median interaction distances are 119 kb and 60 kb, respectively. At these distances, the genomic segments between the anchors often contain unrelated gene(s). The number of spanned genes (Fig. 4C), cautions against the possible assumption that the cCREs typically regulate the neighboring gene. We also counted the number of interactions associated with a single promoter and found that one promoter can interact with up to eight targets (Fig. 4A, D). At lower resolutions, both the number of genes spanned by an interaction and the numbers of significant interactions per promoter (Suppl. Table 3) increase (Suppl. Fig. 7A, B), because low-resolution analyses cannot resolve short-distance and favor longer-distance contacts. The chromosomal distribution of all interactions was skewed towards gene-rich sub-telomeric regions, as expected. Interestingly, the enrichment in H3K27me3 interactions was higher than that of H3K4me3 interactions (Fig. 4E), possibly due to the prevalence of genes regulated by Polycomb in distal chromosomal regions, related to chromosome partitioning^53^.

**Fig. 4.**
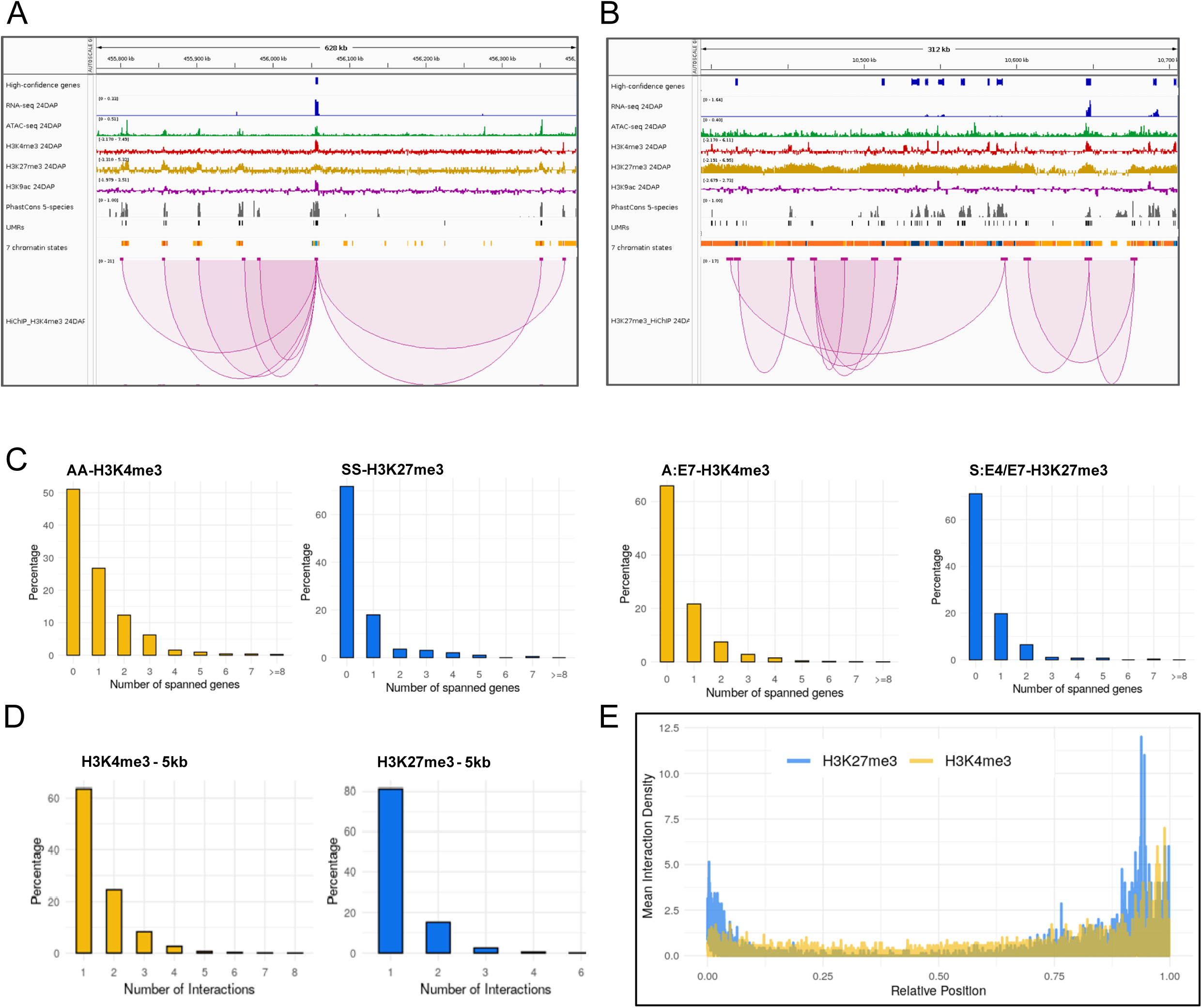
High-resolution analysis of the interactome in the 24DAP embryo. HiChIP data enriched in activating (H3K4me3) or silencing (H3K27me3) histone marks were analyzed at 5- kb resolution. Examples of multiway HiChIP interactions involving (A) an active promoter and (B) a Polycomb-silenced region. (C) Numbers of genes spanned by specific types of interactions: Active promoter-Active promoter (AA), Silent promoter-Silent promoter (SS), Active promoter-E7 state segment (A:E7), and Silent promoter-E4 or E7 segment (S:E4/E7). (D) Numbers of interactions per active promoter (H3K4me3 interactions) and per silent promoter (H3K27me3 interactions). (E) Generalized chromosomal distribution of equal numbers of H3K27me3 and H3K4me3 interactions.

### Annotation of genomic interactions determines the composition of the primary interaction classes and types of involved promoters

To gain a deeper understanding of these interactions and the promoters involved, we hierarchically annotated the interaction datasets. Given that H3K4me3 and H3K27me3 predominantly mark active and silent promoters, respectively, these two genomic features were given top priorities in the annotation, followed by other genic features as terminators and introns. Chromatin states E7 and E4, including cCREs and intergenic Polycomb regions, respectively, ranked further down in the annotation hierarchy, completed with TEs. We identified diverse genomic feature pairs at anchors, defining distinct classes of interactions (Suppl. Fig. 8). Quantification of all possible interaction class proportions (Suppl. Fig. 9A, B) and those involving only active promoters (Fig. 5A, B), revealed that H3K4me3-HiChIP primarily captures interactions between two active promoters (25.8/31.7% for all-interactions and active-promoter-centric analysis, respectively) or active promoter-E7 interactions (22.6/27.8%). Additionally, 7.95% of active promoters interact with introns, and 5.22% with silent promoters. For H3K27me3 silent promoter-centered interactions, E4 and E7 interactions account for 29.4% and 15.3%, respectively, while silent promoter-silent promoter interactions make up 17.6%. We also identified a category of long self-looped genes (Suppl. Fig. 8B), comprising 232 active and 137 silent promoter-terminator pairs within the same gene.

**Fig. 5.**
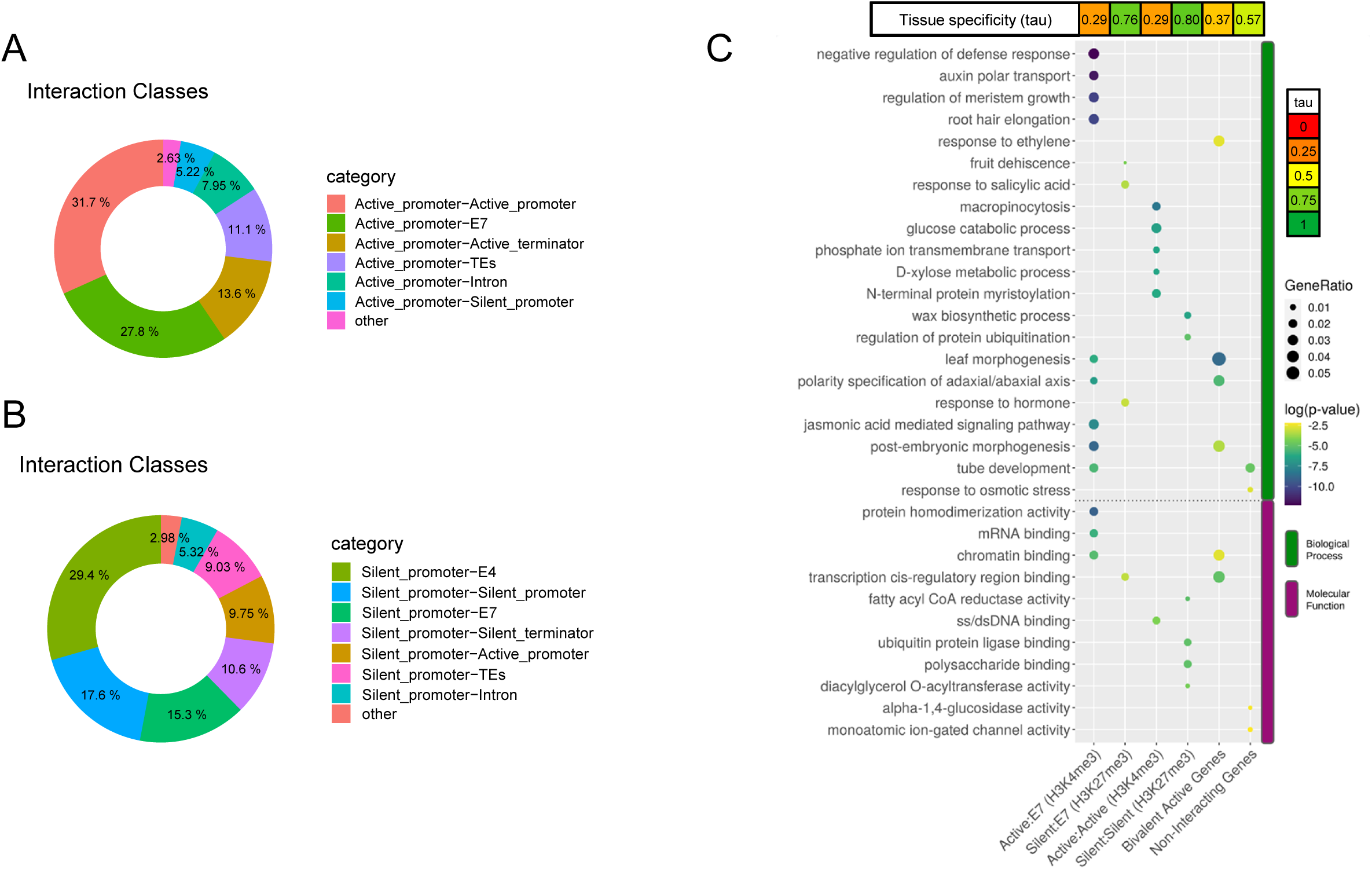
Annotation of HiChIP interactions in the 24DAP embryo. (A) Annotation of significant active promoter-centered (H3K4me3) interaction classes. (B) Annotation of significant silent promoter-centered (H3K27me3) interaction classes. Active promoters, silent promoters, terminators, introns, CRE candidates and transposable elements were used in this order as genomic features for hierarchical annotation. ‘Other’ includes all interactions that do not exceed 5% of the total. (C) GO and tissue specificity analysis of genes involved in specific classes of interactions: Active:E7 (H3K4me3) = Active promoter-E7 genomic segment, H3K4me3 interactions; Silent:E7 (H3K27me3) = Silent promoter-E7 genomic segment, H3K27me3 interactions; Active:Active = Active promoter-Active promoter; Silent:Silent = Silent promoter- Silent promoter.

### Promoter interactions indicate co-regulation or expression enhancement by the interacting partner

Given that about a third of the H3K4me3 interaction set comprises interactions between two active genes (Fig. 5A, Suppl. Fig. 9A), we investigated whether these interactions reflect the physical proximity of genes that are co-regulated during plant development. To test this hypothesis, we assessed whether gene expression changes between stages are more concordant within these interaction pairs than expected by chance. We assigned interacting genes to expression clusters generated by k-means clustering from public gene expression datasets^54,55^ and evaluated the significance of both interaction partner genes falling into the same cluster. Out of 1,988 active promoter-active promoter interacting pairs with a single-gene annotation in each interacting bin, 502 pairs belonged to the same expression cluster. The low p-value (Chi-square test, 4.807e-09) indicates that interacting genes are significantly more likely to belong to the same expression cluster than would be expected by random chance, implying their co-expression and co-regulation. Gene Ontology (GO) analysis suggests that products of these genes play roles in fundamental cellular processes, such as metabolism of monosaccharides and ss/dsDNA binding (Fig. 5C).

We found active genes that interact with extremely low expressed genes or with silent genes. Since these genes were not co-expressed, we asked whether their interactions may serve to enhance the expression of the active gene. We tested whether the expression levels of the active interacting genes exceeded the expected level of an average expressed gene at 24 DAP. The Wilcoxon Rank Sum Test (p-value < 4.618e-06) confirmed that genes interacting with silent promoters of other genes have higher expression values compared to all expressed genes, indicating their upregulation. We hypothesize that these active genes may be enhanced by their interaction with silent promoters or promoter-proximal enhancers.

### Promoter-CRE interactions and bivalent chromatin predominantly involve genes encoding transcription factors

GO annotation of genes involved in active promoter-E7 interactions revealed that this group is dominated by genes encoding proteins involved in morphogenesis, defense response – known to be associated with seed maturation^37^ - and chromatin binding (Fig. 5C) that have low tissue specificity, consistent with the known pleiotropic and multifunctional nature of developmental TFs^56^. To support this observation, we compared genes interacting with E7 segments to a group of non-interacting expressed genes. The first group overlapped significantly more than expected with a set of 2,060 barley developmental transcription factors defined by^55^ (p-value 0.00356, hypergeometric test), whereas the non-interacting group showed no overlap (p-value 0.99). This provides additional evidence that TF genes, in particular, are targets of distal CREs, similar to findings in animals^57^.

Polycomb-silenced genes form loops with other silent promoters as well as E7 and E4 segments. GO term analysis of silent gene-E7 interactions also pointed to TFs (DNA binding *cis-*regulatory activity), however, this set of genes appears highly tissue-specific (values ∼0.8). Additionally, silent promoter-silent promoter H3K27me3 loops were associated with tissue-specific genes, although these genes were clearly linked to metabolic functions. Thus, Polycomb loops may act as important determinants of developmental and tissue-specific regulation.

We further focused on a subset of 355 interactions where both anchors simultaneously overlapped in H3K4me3 and H3K27me3 HiChIP datasets at 5-kb resolution. Interestingly, these interactions involve both active and silent genes. The largest group (97; 27.3%) of these bivalently marked interactions involve active promoters interacting predominantly with the E7 class (Suppl. Fig. 10). This group includes tissue-unspecific genes functionally annotated as TFs, enriched in GO terms such as morphogenesis and chromatin binding (Fig. 5C).

### Late embryogenesis abundant genes interact and are co-regulated

To exemplify the dynamics of the epigenomic landscape and its impact on gene transcription, we focused on a cluster of genes encoding late embryogenesis abundant proteins from the LEA_5 group. This cluster consists of two groups of paralogous genes (Ensembl Plants^58^): *HORVU.MOREX.r3.1HG0061770* (named *LEA*_*A-1*), *HORVU.MOREX.r3.1HG0061780* (*LEA*_*A-2*) and *HORVU.MOREX.r3.1HG0061820* (*LEA*_*A-3*), forming the paralogous group LEA_A, and *HORVU.MOREX.r3.1HG0061790*(*LEA*_*B-1*) and *HORVU.MOREX.r3.1HG0061800* (*LEA*_*B-2*), forming the group LEA_B (Fig. 6). While these genes exhibit negligible or no transcription in most tissues analyzed by^53,55^ (Fig. 6A, B), they show high to ultrahigh transcription levels (630-24,977 TPM) in the maturing embryo at 24 DAP and 32 DAP^55^ (Fig. 6B). Despite these differences, all LEA genes in the cluster share a similar epigenetic landscape during development. Genic and adjacent regions are unmethylated, with minimal differences between 24DAP embryo and leaf tissue. The burst of transcription at 24 DAP is preceded by chromatin opening in promoter regions, which is already apparent in the 8DAP embryo. Later in embryo development, part of the silencing H3K27me3 marks loaded on broader genic regions are replaced by activating H3K4me3 and H3K9ac marks. This process is gradually reverted during seed germination and further development, concurrent with transcription silencing (Fig. 6C). Collectively, the dynamics of epigenomic features across four stages indicate that the ratio of activating versus silencing histone marks in a gene promoter region is the strongest indicator of transcription activity in these developmentally regulated genes.

**Fig. 6.**
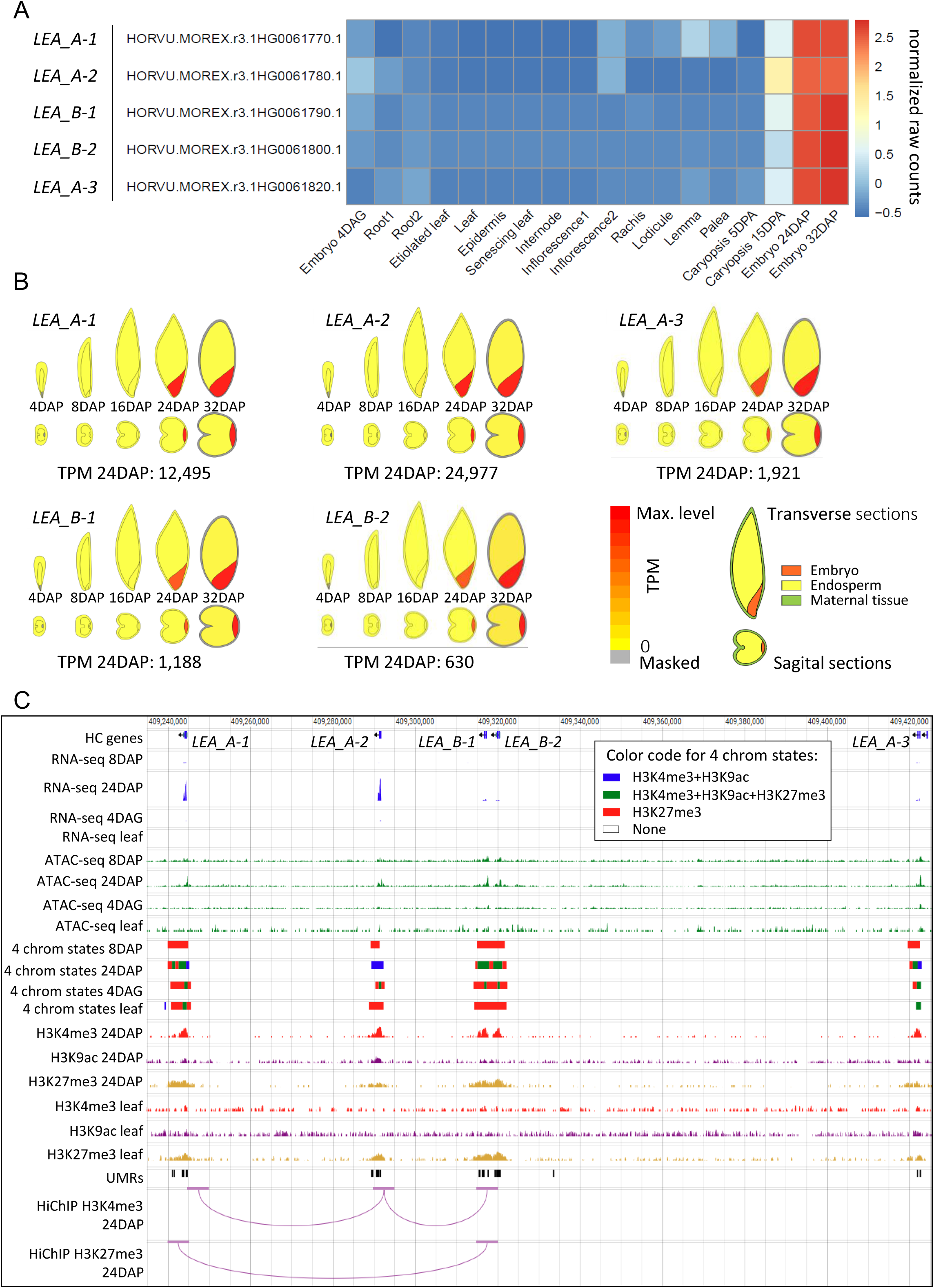
**Transcription, epigenetic landscape, and chromatin interactions in a cluster of barley LEA_5 genes**. (A, B) Results of transcriptomic analysis displayed as (A) differential analysis heatmap showing normalized raw counts for five LEA genes across 18 barley tissues (data from^53,55^), and (B) expression values of the LEA genes across tissues of five stages of a developing barley grain (data from^55^; visualization from barley ePlant database^79^). (C) Visualization of the gene cluster in a genome browser, integrating RNA-seq data^53,55^ and information about open chromatin (ATAC-seq), histone modifications (ChIP-seq and four chromatin states, generated by ChromHMM^5^), and unmethylated regions (UMRs). Activating (H3K4me3) and silencing (H3K27me3) chromatin interactions (HiChIP) were analyzed at 5-kb resolution.

Exploration of activating (H3K4me3) interactions in HiChIP data for the 24DAP embryo indicated that coordination of epigenetic changes, and consequently transcription within the *LEA* cluster, could be mediated by chromatin interactions involving the genes. Analysis at 5-kb resolution revealed highly significant contacts between the centrally located *LEA*_*A-2* gene and its closest neighbors, while no contact was identified between the most distal and the least expressed of the LEA_A paralogs - *LEA*_*A-3* (Fig. 6C). Besides the H3K4me3-mediated contact, the downstream region and/or gene body of *LEA*_*A-1* is connected through an H3K27me3-mediated interaction with the paralog pair *LEA*_*B-1* and *LEA*_*B-2*, confirming spatial contacts of the clustered genes. Interestingly, *LEA*_*A-2*, occupying a central position in the local interactome, has the highest transcription level (24,977 TPM) (Fig. 6B), suggesting a potential positive effect of the spatial organization of the *LEA* locus on the gene’s transcription. This hypothesis is supported by the fact that promoter regions of *LEA_A-2* interactors - *LEA_A-1*, *LEA_B-1* and, possibly, *LEA_B-2* -contain binding sites for ABI5 homolog (*HORVU.MOREX.r3.3HG0300770*, motif ACGTGTC), a bZIP transcription factor known to regulate LEA genes^37^, while no such site was identified by the PlantTFDB prediction tool within 1,500 bp upstream of the TSS of *LEA A-2*, as identified by CAGE^5^. Experimental validation will be required to confirm the potential collaboration of these promoters in transcription regulation.

### Regulatory elements of *Vrn3* are detectable in non-expressing tissues and without vernalization treatment

Finally, we used information about known enhancers of the bread wheat *Vrn3* gene^41^ to test the predictive power of our datasets. The wheat enhancers, located within 30 kb upstream of *Vrn3*, were identified through differential analysis of open chromatin after vernalization treatment of winter wheat. Here, we used spring barley cv. Morex, which does not require vernalization treatment to flower, and explored the epigenetic landscape and interactome in the upstream region of barley *Vrn3* ortholog, *HORVU.MOREX.r3.7HG0653910*. Despite using embryonal and leaf tissues, with no (embryo) or minimal (leaf) transcription, we were able to predict four cCRE regions, showing an overlap of evolutionary conserved sequences with chromatin features corresponding to active or silenced CREs, within 330 kb upstream of the gene (Fig. 7A). A cCRE located 242 kb upstream of *Vrn3*, designated cCRE3, exhibited a high-confidence (FDR < 0.05, 5-kb resolution) H3K27me3-associated interaction with the gene. A BLAST search for the published wheat enhancers (P3 and P4)^41^ provided hits overlapping with barley cCRE3 and cCRE4, respectively (Fig. 7B, C). These CRE candidates are characterized by a bivalent epigenetic state, just like the *Vrn3* promoter, suggesting that this developmental gene is primed but remains silenced until it receives the final trigger for expression, likely through the age-related pathway involving SPL7/15^41^.

**Fig. 7.**
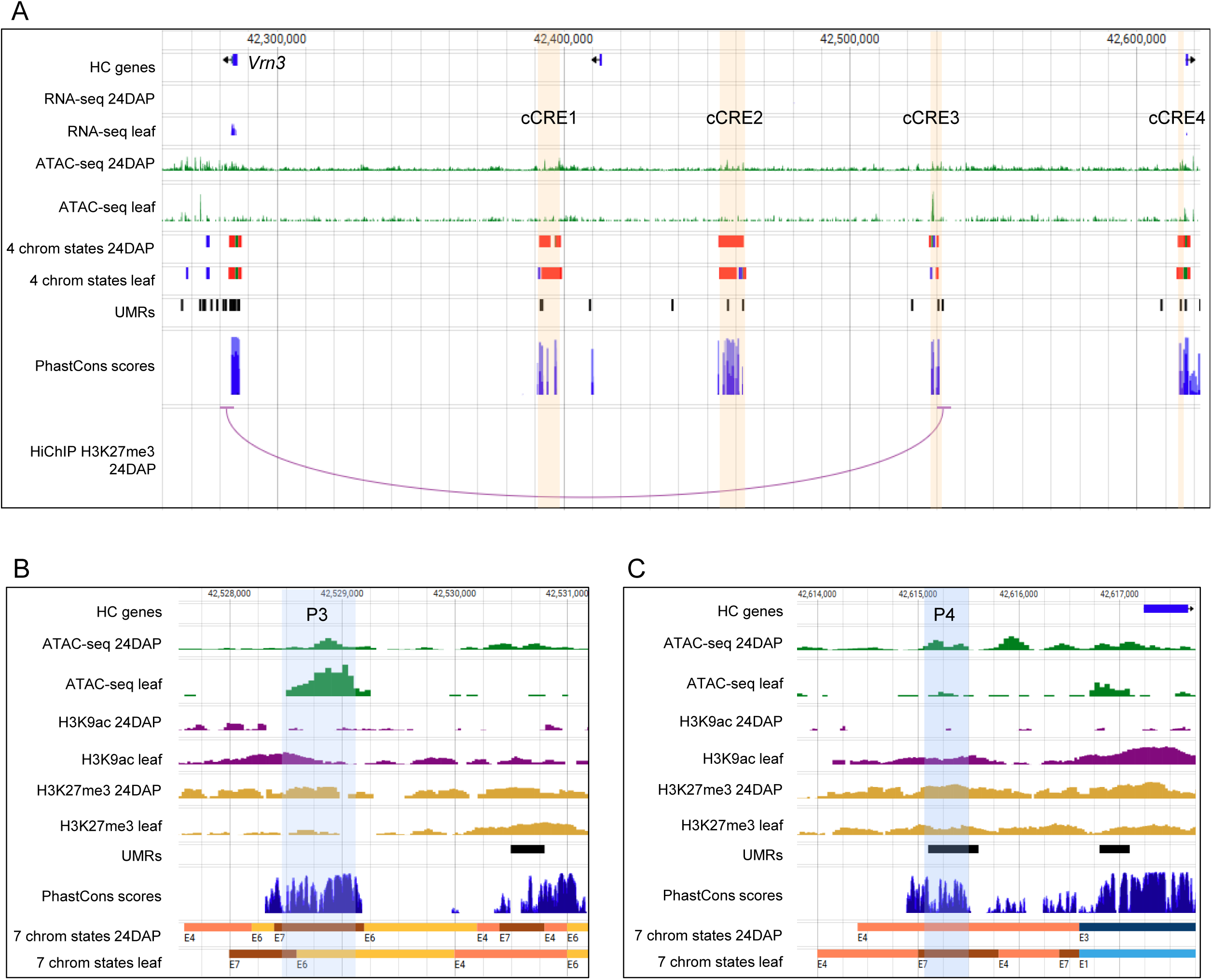
Regulome of the barley *Vernalization 3* gene. (A) Barley *Vrn3* (*HORVU.MOREX.r3.7HG0653910*) locus with cCRE regions (orange bars), predicted based on an overlap of evolutionary conserved sequences (PhastCons scores) with chromatin features associated with active or silenced CREs. Note the long-range (242-kb) contact between *Vrn3* and the cCRE3 region. The color code for four chromatin states is the same as in Fig. 6. (B, C) Zoom-in views of the cCRE3 (B) and cCRE4 (C) regions with BLAST hits of bread wheat P3 and P4 enhancers^41^ (blue bars).

### Dataset visualization in public databases enables application in barley research

Our comprehensive analysis of barley epigenomic landscapes and chromatin interactions highlights the functional significance of CREs in regulating transcription during development. The resulting barley CRE and interactome collections, together with profiles of several epigenetic marks and evolutionary conservation, complemented by published transcriptomic data^5,53,55^, all based on MorexV3 genome^59^, serves as a valuable resource for other researchers. To ensure accessibility, we will make all processed data files and their visualizations available in a JBrowse interface through the Elixir platform (https://olomouc.ueb.cas.cz/en/resources/barleyepibase).

## Discussion

We integrated the most informative epigenetic features using an overlap-based and machine-learning approach, demonstrating that while peak overlap defines a robust cCRE set, the ChromHMM approach offers a broader, more informative classification,b overcoming dataset variability limitations^60^. To maximize cCRE identification, while maintaining analytical simplicity, we selected a 7-state ChromHMM model, which identified 1.43% of the barley genome as having regulatory potential (E7 chromatin state), consistent with 1.5% reported for wheat seedlings^7^. The H3K27me3 E4 state, reflecting intergenic Polycomb-mediated silencing^27^, may contain inactive CREs often escaping detection, manifested as small UMRs embedded in the E4 blocks.

The occasional presence of H3K27me3 in E7 (Fig. 1C) indicates that E7 segments may include not only active cCREs, but also elements in transitional or binary states, as well as those active in some cell types but repressed in others within the sample. Benchmarking chromatin state revealed that E7 could be further divided into sub-states, providing deeper insight into regulatory element dynamics. While E7 dynamics across the four barley stages may appear limited (Fig. 2A), the large number of commonly detected elements aligns with the number of commonly active genes in transcriptome analyses, reflecting pleiotropy and combinatorial dependence of enhancers^61,62^.

The largest comparative study of unstable RNA across diverse plants and vertebrates by capped small RNA sequencing^34^ found that unstable and bidirectional transcription is rare in plants. In contrast, Xie et al.^35^ detected active transcription of thousands of enhancers in the bread wheat genome using two independent techniques (pNET-seq and GRO-seq). Their observation that genes associated with transcribed enhancers are expressed at significantly higher levels lead them to elevating the predictive power of eRNA above other chromatin features. Using a yet-different technique, NET-CAGE, we failed to detect many bidirectionally transcribed cCREs or upstream antisense RNAs, common in animals. Still, a small subset of robust cCREs and E7 elements overlapped with unbalanced or unidirectional nascent capped RNA. Interestingly, transcripts belonging to cluster C4 (Fig. 4F) initiated from low-complexity GA-microsatellite regions, which are described in wheat as BPC5/Ramosa TFBS and are overrepresented in wheat distal CREs^35^. Overall, barley had far fewer transcribed cCREs than found in wheat, suggesting that enhancer transcription plays a minor regulatory role in this species, consistent with findings of McDonald et al.^34^ While methodological differences among studies might contribute to these discrepancies^63^, they likely reflect fundamental differences in transcriptional regulation mechanisms between plants and animals^9^ worth further exploration.

Previous *cis-*regulation studies without 3D structural assays often inferred CRE targets based on proximity or expression levels of neighboring genes. However, our counting of genes interposed between interacting partners demonstrates that this assumption can lead to false conclusions. Also gene expression level alone may reflect the gene promoter activity rather than upregulation by CRE. 3C-based techniques can discern enhancer-promoter interactions but require high resolution, which is challenging to reach in complex tissues and repeat-rich genomes with low mappability, such as that of barley. Our analysis at 5-kb resolution hindered precise annotation of gene-proximal and intronic CREs. While hundreds of intronic CREs were observed in our study, consistent with findings in the human genome^64^, they are not presented due to potential gene-type bias. The resolution also affects interaction distances. The median distance of H3K4me3 barley HiChIP loops at 20-kb resolution (220 kb) is much greater than that at 5 kb (60 kb), but is consistent with results from wheat low-resolution RNA polymerase II- HiChIP (200-400 kb)^20^. Notably, the long-distance interactions may reflect structural rather than functional loops, akin to topologically associating domains.

Our HiChIP data identified bivalent interactions of chromatin segments simultaneously marked as both activating (H3K4me3) and repressive (H3K27me3), typically in developmental regulatory genes that are poised for activation or silencing^65^. In plants, evidence of bivalent chromatin is emerging in both developmental and stress-responsive genes^66,67^. Our GO and tissue-specificity analyses of interacting genes (Fig. 5C) indicated that H3K27me3 silences genes for tissue-specific transcription factors via their interactions with intergenic Polycomb regions, while pleiotropic developmental transcription factors engage in binary interactions. However, distinguishing true bivalent chromatin, with both marks on the same nucleosome, from allelic or sample heterogeneity remains challenging, requiring sequential ChIP for confirmation^68^.

We unveiled a rich promoter-promoter interactome, which is indicative of co-expression hubs and transcription factories, reported in other plant species^20,69,70^. Using pairs of active and lowly expressed or silent genes, we explored the hypothesis that promoters can function as enhancers, and found support for this phenomenon in barley. While this could stem from insufficient HiChIP resolution, failing to discern core promoters from proximal CREs, it is consistent with reports of promoters with regulatory potential both in mammals^71^ and plants^34^. Zhu et al.^69^ proposed a model of active transcription hubs that unifies the roles of active promoters and enhancers, assigning general enhancer-like functions to active promoters, which was also supported by other studies^72,73^. Our exploration of TFBSs in promoters of interacting LEA genes lead us to a hypothesis that the ultrahigh transcription of *LEA_A-2* gene, positioned centrally in the local interactome, might be due to the enhancer-like function of the interacting promoters, bringing ABI5 TFs, whose BS is missing in *LEA_A-2* promoter. Alternatively, the promoter of *LEA_A-2* may be the core regulatory element in the gene cluster, supporting transcription of its interacting partners. In this scenario, the trigger of the transcription burst would not be ABI5 but another upstream regulator. Analysis of structural variation in promoter regions of the interacting *LEA* genes across multiple genotypes and their correlation with transcription levels may shed light on this phenomenon.

Our investigation of the *Vrn3* region demonstrated the potential of our datasets to show signatures of distal CREs even in non- or low-expressing tissues and without the environmental stimulus that led to their identification in winter wheat^41^. It also indicated a relationship of these enhancers and the age-related pathway in spring barley. A silencing long-range interaction in barley *Vrn3* region revealed a spatial contact between one of our cCREs and *Vrn3*, particularly remarkable given the tenfold greater distance between them in barley than in wheat. A study using ChIA-PET for H3K4me3^42^ found no significant chromatin loop in the *Vrn3* region after vernalization in winter wheat. The authors suggested that this might reflect higher enrichment in H3K27me3 than H3K4me3 in the locus, which aligns with our findings and confirms the region’s bivalent chromatin state.

To validate CRE candidates, large-scale enhancer activity assays such as MPRA or Plant STARR-seq^74,75^ could be used to assess enhancer functionality and promoter-enhancer compatibility. However, testing elements outside their genomic context may yield inconclusive results, necessitating targeted editing within the native loci for definitive validation. Since transcriptional changes drive a range of advantageous traits, CREs hold significant promise for trait engineering. Advances in machine learning, neural networks, and applications of large language models^76,77^ enable to leverage high-quality datasets, such as those generated here, for model training. These tools enhance understanding of plant regulomes and support their biotechnological applications^78^.

## Resource availability

### Lead contact

Requests for further information and resources should be directed to and will be fulfilled by the lead contact, Hana Simkova.

### Data and code availability

Previously generated ChIP-seq datasets are available in the GEO database under accession number GSE227218. The datasets generated during the current study include outputs from ATAC-seq, ChIP-seq (leaf), NET-CAGE-seq, HiChIP, and bisulfite-seq (24DAP embryo). The data were deposited in the Sequence Read Archive (SRA) - PRJNA1177611. All processed datasets will be made available for download and visualization in the Jbrowse interface at https://olomouc.ueb.cas.cz/en/resources/barleyepibase.

Supplemental data and high-resolution figures are deposited in GitHub repository https://github.com/MorexV3CAGE/Barley_distal_regulome.

This paper does not report original code.

## Acknowledgments

The project was supported by the Czech Science Foundation (grant number 21-18794S) and from the project TowArds Next GENeration Crops, reg. no. CZ.02.01.01/00/22_008/0004581 of the ERDF Programme Johannes Amos Comenius. Computational resources were provided by the e-INFRA CZ project (ID:90254), supported by the Ministry of Education, Youth and Sports of the Czech Republic. We thank Jitka Weiserova for technical assistance in flow sorting, Zdenka Bursova for plant maintenance, Katerina Holusova, Helena Tvardikova, Pascal Jaroschinsky, Axel Himmelbach, Jörg Fuchs for their assistance with library preparation and sequencing. The work performed at IPK Gatersleben was part of the p-epBAR project, supported by the German Ministry of Education and Research (BMBF) (grant number FKZ 031B1224). Jbrowse genomic browser is provided by ELIXIR-CZ Research Infrastructure Project [LM2023055].

## Author contributions

HS, PNa, NS: Project conceptualization; PNa, SP, ZT, ZZ: Data acquisition: PNa, SP, OK, HS, PNo, ZT: Formal analysis and data management; PNa, SP, HS: Writing - Original Draft; PNa, HS, SP, ZZ: Writing; HS, PNa, ZZ, NS: Review & Editing; HS, NS: Funding acquisition.

## Declaration of interests

The authors declare no competing interests.

## Declaration of generative AI and AI-assisted technologies

During the preparation of this work, the authors used ChatGPT4o in order to refine English language and remove possible redundancies, without changing the information content and meaning. After using this tool or service, the authors reviewed and edited the content as needed and take full responsibility for the content of the publication.

## Supplemental information titles and legends

**Supplemental_Figures.** Suppl. Figures 1-10

**SupplFile_1_Coordinates of E7 segments in MorexV3 genome** – across four stages

**SupplTable_1_PeakNumbers** – peak numbers in barley embryo ChIP-seq and ATAC-seq datasets

SupplTable_2_NET-CAGE_statistics_and_quality_control

**SupplTable_3_Interaction numbers** - numbers of interactions in all HiChIP samples as identified by FitHiChIP

## STAR Methods

### EXPERIMENTAL MODEL DETAILS

#### Plant material

Barley cv. Morex was grown in growth chambers (Weiss Gallenkamp, walk-in chambers) at 16/8 hrs light cycle, 16 °C day/12 °C night temperature. For the 4DAG seedlings, seeds were germinated on wet tissue paper at 20 °C for 4 days before harvesting and removing remnants of seed coat and endosperm. The 8- and 24DAP embryos were staged according to their time of fertilization, size and phenotype and dissected as described previously^80^. For collecting leaf samples, plants were grown in growth chambers at 16/8 hrs light cycle, 20 °C day/16 °C night temperature for two weeks.

### METHOD DETAILS

#### ATAC-seq

Barley embryos were fixed for 8 min in 1% methanol-free formaldehyde (Pierce^TM^ 28906) in PBS under vacuum. Fixation was stopped by 5 min incubation in 0.125M glycine in PBS followed by thorough PBS washes. The tissues were pulverised by mortar and pestle in liquid nitrogen and nuclei were extracted in lysis buffer LB01^81^ supplemented with cOmplete™, Mini, EDTA-free protease inhibitor (Roche). Twenty-five thousand G1 nuclei per sample, counterstained with DAPI, were purified by FASCAria II SORP flow cytometer and sorter (BD Bioscience, San Jose, USA) and processed using the ATAC-seq kit (Active Motif 53150) with a decrosslinking step included before DNA isolation. Tagmented and amplified libraries were sequenced on the NovaSeq6000 platform (Illumina).

Fresh leaf samples from two-week-old seedlings were finely chopped using a razor blade in nuclei isolation buffer (0.25 M sucrose, 10 mM Tris-HCl pH 8.0, 10 mM MgCl2, 1% Triton X-100, 5 mM β-Mercaptoethanol) supplemented with 1x Halt™ Protease Inhibitor Cocktail (Thermo Scientific). Seventy-five thousand nuclei per sample were estimated by flow cytometry sorting and processed using Tagment DNA Enzyme (TDE1, Illumina, 20034197). Transposition products were purified using the MinElute PCR Purification Kit (QIAGEN, 28004), amplified using the NEBNext® High-Fidelity 2x PCR Master Mix (NEB, M0541), and further purified using the VAHTSTM DNA Clean Beads (Vazyme, N411). The final libraries were sequenced on the Illumina NovaSeq 6000 platform (Illumina) at IPK Gatersleben.

#### ChIP-seq

The ChIP experiment for leaf was performed according to a previously described protocol^82^with minor modifications. Fresh leaves (3 g) from two-week-old seedlings were fixed under vacuum in 1% formaldehyde (Sigma-Aldrich 252549) for 15 min. Fixation was stopped by a 5-min incubation in 0.125 M glycine. The tissues were pulverised using a mortar and pestle in liquid nitrogen and nuclei were extracted. Nuclei samples were resuspended in Nuclei Lysis Buffer containing 0.1% SDS in a 1 ml sonication tube (Covaris). Chromatin was sonicated for 250 s in a Covaris S220 with settings of peak power 175 W, cycles/burst 200, and duty factor 20%. The sonicated chromatin was cleaned by centrifugation, and the supernatant was diluted four times using ChIP Dilution Buffer. The diluted chromatin (800 µl) was incubated with the respective antibodies (anti-H3K4me3, Abcam 213224; anti-H3K27me3, Abcam 6002; anti-H3K9ac, Abcam 4441) at 4°C for 16 h. Washed Dynabeads™ Protein A (Invitrogen), 40 µl per sample, were added to the antibody-bound chromatin and incubated at 4°C for 2 h. The collected beads were washed twice sequentially in Low Salt Buffer, High Salt Buffer, and TE Buffer. The bead-bound chromatin was purified using the iPure kit (v2, Diagenode C03010015) according to the manufacturer’s instructions. Purified DNA was quantified using the Qubit^TM^ dsDNA High Sensitivity Assay kit (Invitrogen). ChIP-seq libraries were prepared using the NEBNext® Ultra™ II DNA Library Prep Kit for Illumina (New England Biolabs) and sequenced on the Illumina NovaSeq 6000 platform (Illumina) at IPK Gatersleben.

#### Bisulfite sequencing

DNA for preparation of whole genome bisulfite sequencing (WGBS) libraries was isolated from three frozen 24DAP embryos using NucleoSpin Plant II kit (Macherey-Nagel) in two biological replicates. DNA was quantified using a Qubit high-sensitivity DNA kit (Thermo Fisher Scientific). To assess the conversion efficiency, 100 ng input DNA was spiked with 1 ng of *E. coli* DNA. Both bisulfite conversion and WGBS library preparation were done using a Zymo-Seq WGBS Library kit (Zymo Research) following the manufacturer’s instruction with one modification, namely that a tagmentation step was shortened from 15 to 10 minutes. Paired-end 2x150bp reads were generated using the Illumina NovaSeq 6000 platform.

#### HiChIP

The 24DAP embryos were fixed under vacuum in 2% formaldehyde in PBS for 15 min. Fixation was stopped by 5 min incubation in 0.125M glycine in PBS and subsequent PBS washes. The tissues were pulverised by mortar and pestle in liquid nitrogen and nuclei were extracted in lysis buffer LB01^81^ supplemented with cOmplete™, Mini, EDTA-free protease inhibitor (Roche). Five million G1 nuclei were sorted by FASCAria II SORP flow cytometer into LB01 supplemented with cOmplete™, Mini, EDTA-free protease inhibitor (Roche) to perform digestion and proximal ligation using Arima HiC+kit (Arima Genomics, protocol A101020). HiChIP was performed using Arima HiChIP protocol A160168 v00, followed by library preparation using Accel-NGS 2S Plus DNA Library Kit (Swift Biosciences) according to protocol A160169 v00. For histone-modification enrichment, we used the H3K4me3 (04-745) antibody from Millipore and H3K27me3 (C15410195) from Diagenode.

#### NET-CAGE

Two replicates of 4DAG embryos were collected in liquid nitrogen and grinded by mortar and pestle for nuclei isolation. To avoid transcriptional run-on, we added α-amanitin (a potent inhibitor of transcription) and RNAse inhibitor at each experimental step. We purified nuclei from 4DAG embryos by flow cytometry, followed by isolation of small RNA using PureLink™ miRNA Isolation Kit (Invitrogen). The cap-trapping, library preparation and sequencing were performed in DNAFORM company. The libraries were sequenced on the Novaseq6000 platform (Illumina) at 150bp PE.

### QUANTIFICATION AND STATISTICAL ANALYSIS

#### ATAC-seq and ChIP-seq data analysis

All sequencing datasets were trimmed with Trim Galore (version 0.6.4). The ATAC-seq data was analyzed according to^83^. The ATAC-seq and ChIP-seq reads were mapped (bowtie2 v2.4.2) and normalized coverages were calculated using deeptools bamCoverage and bamCompare (v3.5.1), respectively. MACS2 (version 2.2.7.1) program^84^ with parameters – broad –nomodel was used to identify the peaks.

#### Bisulfite-sequencing data analysis

The bisulfite sequencing reads were trimmed using Trim Galore (v0.6.4) with –trim1 –paired options. Trimmed reads were aligned by the Bismark program (version 0.19.0)^85^ with the default settings, deduplicated and data for each methylation context were generated. Subsequently, we defined intergenic unmethylated regions (UMRs) using the approach described by^13^. We only considered cytosines with minimal coverage of 5 and defined the UMRs by setting a 1% methylation threshold in every context throughout 300bp intergenic fragments. Using publicly available bisulfite sequencing data from leaf tissue^59^, we defined leaf UMRs using the same criteria.

#### HiChIP data analysis

Each replicate contained approximately 500 million reads, which were trimmed and processed by the HiC-Pro (v3.0.0) pipeline^86^. Replicates were merged after the deduplication step for further valid-pair processing. The resulting intra-chromosomal contact maps were further processed by FitHiChIP (Release 6.0)^52^ for significant loop calling using previously generated ChIP-seq peaks as a reference. In this process, 2 Mb distance was set as a maximum distance, nearby interactions were merged, peak-to-all interactions were considered and the FDR value limit was set to 0.05 or 0.1. GenomicInteractions package^87^ was used for the genomic feature annotation of the loop anchors.

#### RNA-seq data processing

Trimmed RNA-seq reads from 8DAP, 24DAP, and 4DAG embryos, as well as leaf tissue, were mapped to the MorexV3 genome using STAR (v2.7.6a), followed by transcript quantification by RSEM (v1.3.3) software package. Genomic coverages were calculated using deeptools bamCoverage (v3.5.1), for visualization. To account for potentially unannotated genes, we have merged all the mapped RNA-seq data into a single bed file using bedtools bamtobed and selected for intergenic regions that had TPM > 1. The lncRNA^45^ coordinates were converted by liftoff software from MorexV1 to MorexV3 assembly. Previously generated expression data^53,55^ were analyzed by DESeq2^88^ for differential expression. For visualization, the data were normalized using the vst function.

#### NET-CAGE data analysis

Sequencing reads were filtered to remove rRNA reads and mapped using BWA with MAPQ<=20 and HiSat2 on MorexV3 reference genome and transcriptome.

Data from both replicas were processed and merged in the CAGEr pipeline (v2.6.1)^89^ with the following settings: (removeFirstG = TRUE, correctSystematicG = FALSE, T = 1mil, alpha = 1.33, TPM threshold = 0.1, TPM singletons to filter < 5). Regions corresponding to chrUn and rDNA were masked from the MorexV3 assembly and, for enhancer analysis, only distal intergenic regions were analyzed. Intergenic regions were defined by ChIPseeker (v1.36.0^90^) annotation (> 500 bp from annotated gene).

Based on our intergenic RNAseq data and lncRNA annotation, the set was further pruned for potential unannotated genes and lncRNAs by removing all the regions close to a significant RNAseq signal (within 500-bp distance of the region, > 1 TPM). Based on previous data from mammals regarding enhancer bi-directionality, the NET-CAGE clusters were split further into unidirectional and bidirectional balanced and unbalanced categories. Clusters that had an antisense NET-CAGE signal of TPM > 0.1 and not singleton in 600-bp neighborhood were considered bidirectional and their size was adjusted to span region from the plus strand- dominant TSS to the minus strand-dominant TSS. The limits for balanced signals were set to log2FoldChange from 1 to -1. Association of the cCREs with NET-CTSSs was tested by a permutation test with 500 cycles using regioneR package^91^.

#### Sequence conservation analysis

Sequences of H. vulgare MorexV3^59^ were used as the reference for pairwise alignments with the sequences of *T. urartu* v2.0^92^, *S. cereale* Rye_Lo7_2018_v1^93^, *B. distachyon* v3.0^94^; International Brachypodium Initiative, 2010), and *Ae. tauschii* v4.0^95^ using aligner AnchorWave (v1.2.5)^96^, followed by processing through the pipeline: axtChain, chainNet, netToAxt, and axtToMaf (UCSC Genome Browser Toolkit, Anaconda distribution). The resulting pairwise alignments were then combined into a multiple sequence alignment (MSA) using aligner ROAST (v3)^102^. PhastCons score was computed using the PhastCons program from the PHAST package (v1.6)^97^. To obtain the neutral and conserved models for PhastCons, the phyloFit program (a part of the PHAST package) was used according to the manual (http://compgen.cshl.edu/phast/phastCons-HOWTO.html). Conservation scores for sets of sequences were calculated as averages from all base pair PhastCons scores included in each sequence or a bin^59^.

#### ChromHMM chromatin state analysis

ChromHMM (v1.23)^46^ was used to learn seven chromatin states for each of the developmental stages. Previously published (embryonic stages^5^) and newly generated (leaf) ChIP-seq and ATAC-seq mapped and deduplicated data were binarized from BAM files, while the ‘coding potential’ and UMRs were binarized from BED files using the ‘-peaks’ option, all with a bin size of 200 bp. Binarized data were used to learn seven chromatin states, followed by Overlap Enrichment analysis. To provide condensed information on the three analyzed histone modifications, leaf ChIP-seq data from^6^ and this study were used alone to learn four chromatin states, complementing similar datasets previously generated for embryonic tissues^5^.

#### RSAT motif analysis

To determine core motifs, sets of cCREs were subjected to peak-motif position-analysis by Regulatory Sequence Analysis Tools (RSAT) integrated with footprintDB^98,99^ and followed by motif-clustering to eliminate redundancy. Custom settings are documented in the rsat_analysis.txt deposited in the GitHub repository.

#### GO analysis

Using our previously generated GOMAP GO annotation^5,100^ and the ‘enricher’ function from the ‘clusterProfiler’ package with default settings, we determined the enrichment of GO terms for interacting and non-interacting genes. To reduce the redundancy of the results we utilized the REVIGO RESTful API^101^. From this pruned set, only the upper quartile of enriched GO terms was used for visualization.

## Notes

https://github.com/MorexV3CAGE/Barley_distal_regulome

### Competing Interest Statement

The authors have declared no competing interest.

