## Supplementary material for "Epigenome and interactome profiling uncovers principles of distal regulation in the barley genome": Suppl. Figures 1-10

A B C

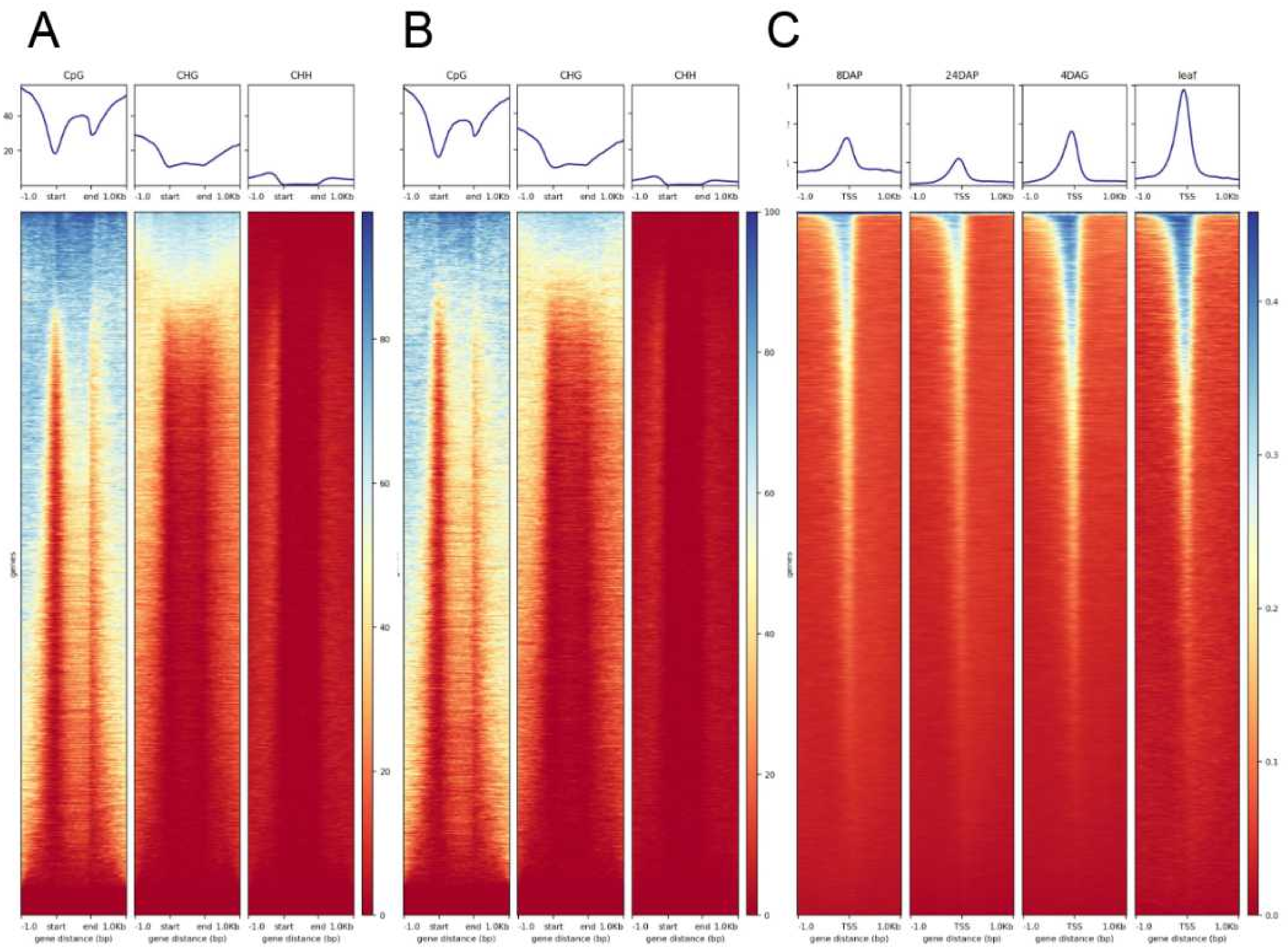


**Suppl. Fig. 1. Profiles and heatmaps showing read coverages for two epigenetic features across MorexV3 high-confidence genes.** Data from bisulfite sequencing show cytosine methylation in three sequence contexts in (A) 24DAP embryo and (B) leaf. (C) Profiles of ATAC-seq coverages around transcription start sites (TSSs) in the four developmental stages used in the study.

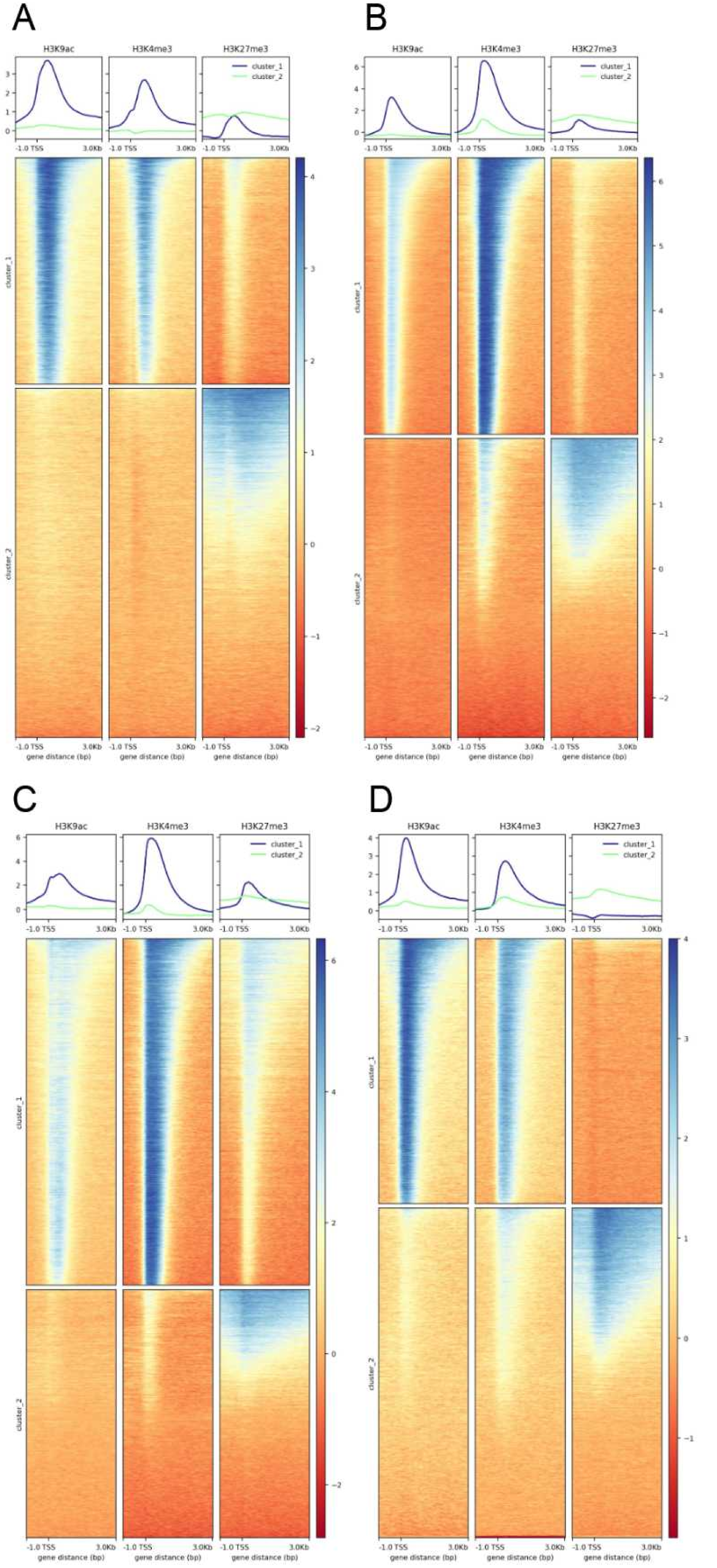


**Suppl. Fig 2**. **K-means clustered profiles of histone modifications around HC-gene TSSs from ChIP-seq**. (A) Profiles in 8DAP embryo. Cluster 1 (top) represents active genes while cluster 2 (bottom) shows profiles of silent genes. (B, C, D) Data from 24DAP (B), 4DAG (C) and leaf (D) samples. The k-means clustering separates two sub-profiles reflecting transcriptional activity.

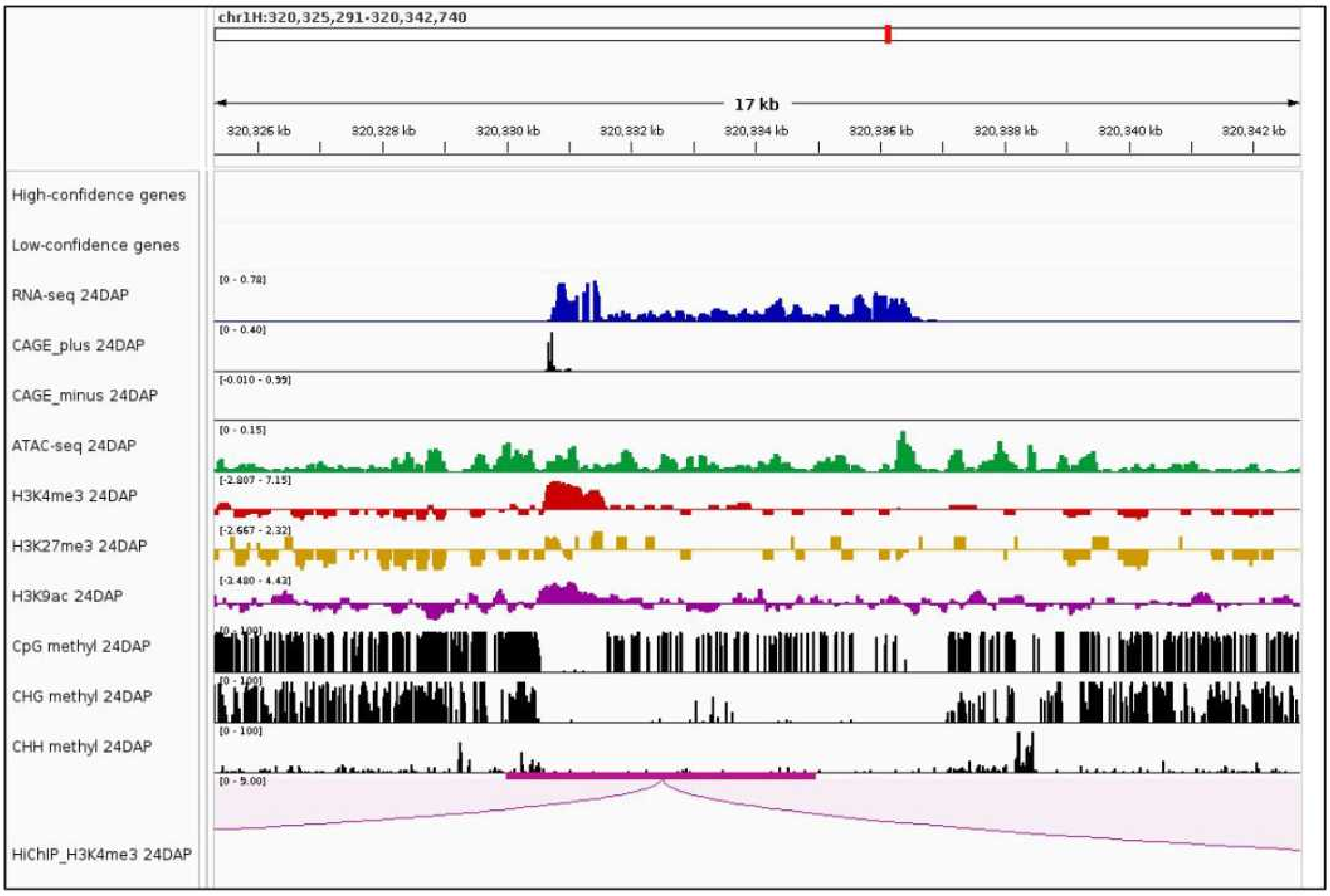


**Suppl. Fig. 3. An example of a putative unannotated gene.** The occurrence of transcripts and features of active chromatin indicates the presence of an unannotated gene. To prevent misinterpretation of its promoter as an intergenic CRE, we excluded this and similar regions from the analysis of the non-coding genome.

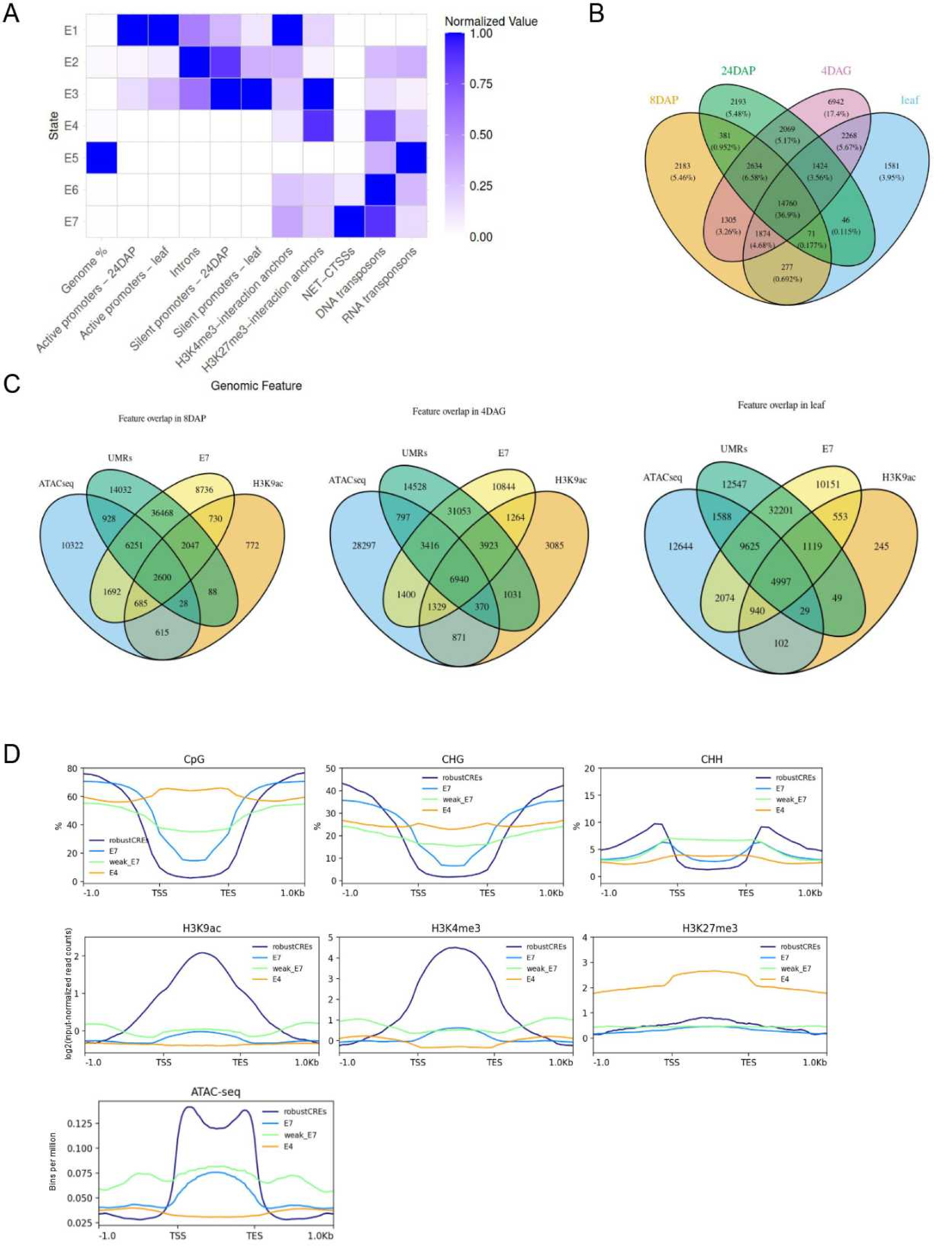


**Suppl. Fig. 4**. **Chromatin state analysis is a sensitive method of cCREs detection.** (A) An overlap enrichment of genomic features across 7 chromatin states in the leaf sample showing state dynamics following transcription dynamics between individual stages. (B) Dynamics of chromatin state-E4 segments, potentially comprising silenced cCREs. (C) Overlaps of the intergenic epigenetic features, as detected by ChIP-seq/ATAC-seq peak calling and DNA methylation analysis, with the E7 chromatin state in 8DAP embryo, 4DAG embryo, and leaf. (D) Comparison of main chromatin feature profiles across robust cCREs, states E4, E7 and the E7-state segments not overlapped with any of the epigenetic feature peaks (weak E7).

A

| 8DAP robust CRE motif | e-value | Transcription factor |
|---|---|---|
| 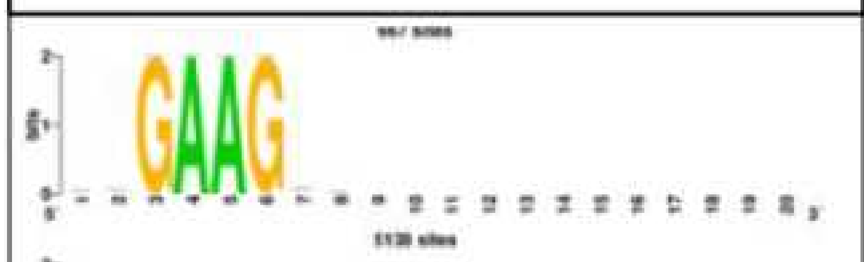 | | GAGA-binding protein like family, BPC1 |
| 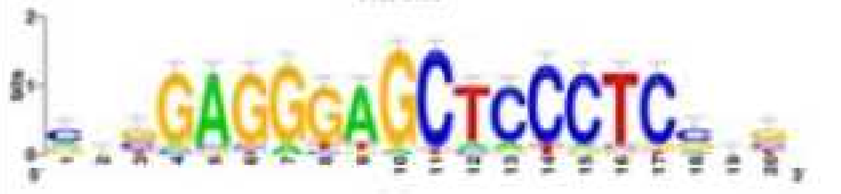 | 6.5e-20 | Palindrome of the GAGA motif |
| 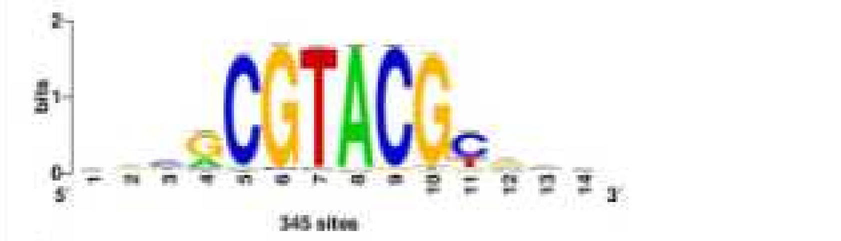 | 5.1e-07 | Squamosa promoter-binding-like protein |
| 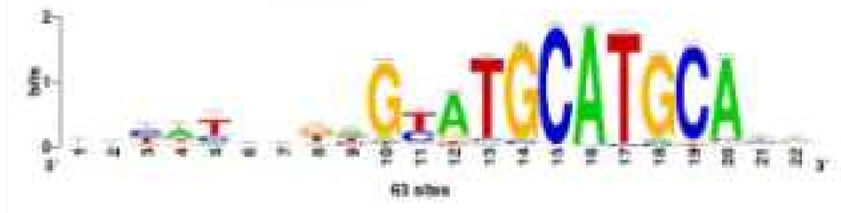 | 3.6e-06 | B3 family protein |

B

| 4DAG robust CRE motif | e-value | Transcription factor |
|---|---|---|
| 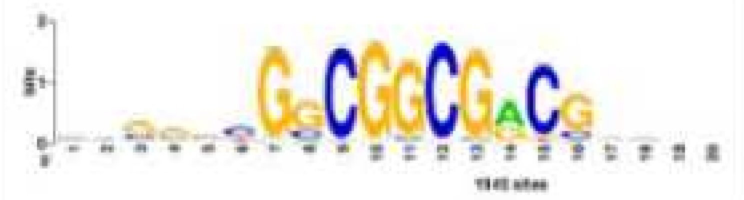 | 7.6e-101 | ERF family protein |
| 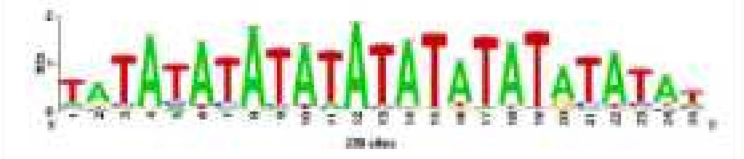 | 1.9e-63 | ZFHD10-3 |
| 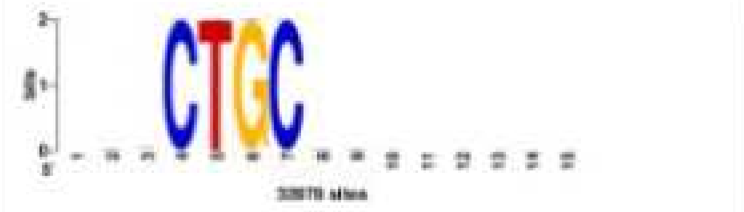 | 9.8e-07 | NAC family, ONAC |
| 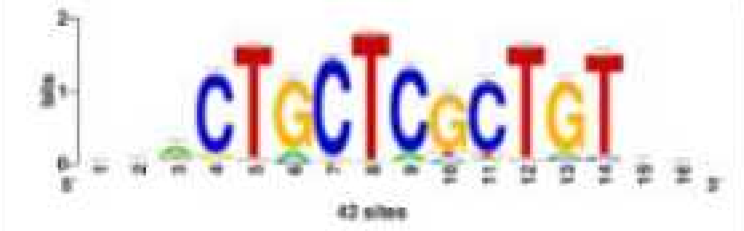 | 0 | No hit |

C

| leaf robust CRE motif | e-value | Transcription factor |
|---|---|---|
| 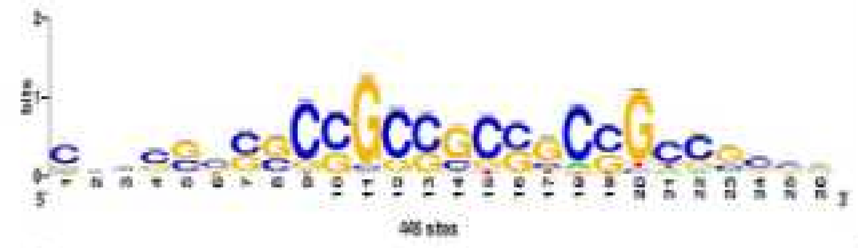 | 1e-300 | ERF family protein |
| 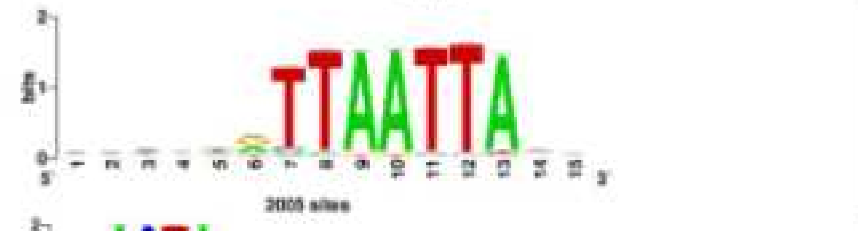 | 1.5e-32 | WOX (Wuschel-like homeobOX) |
| 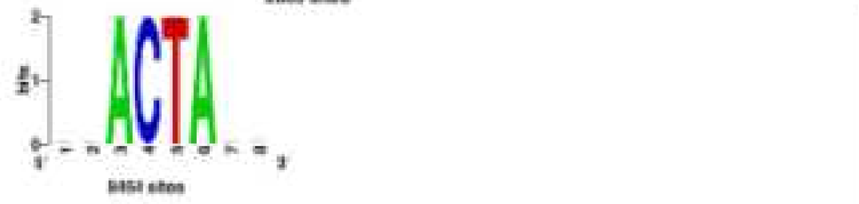 | 8.9e-04 | AP2/ERF and B3 domain-containing protein |

**Suppl. Fig. 5**. Transcription factor binding sites enriched in (A) 8DAP, (B) 4DAG, and (C) leaf robust cCRE datasets.

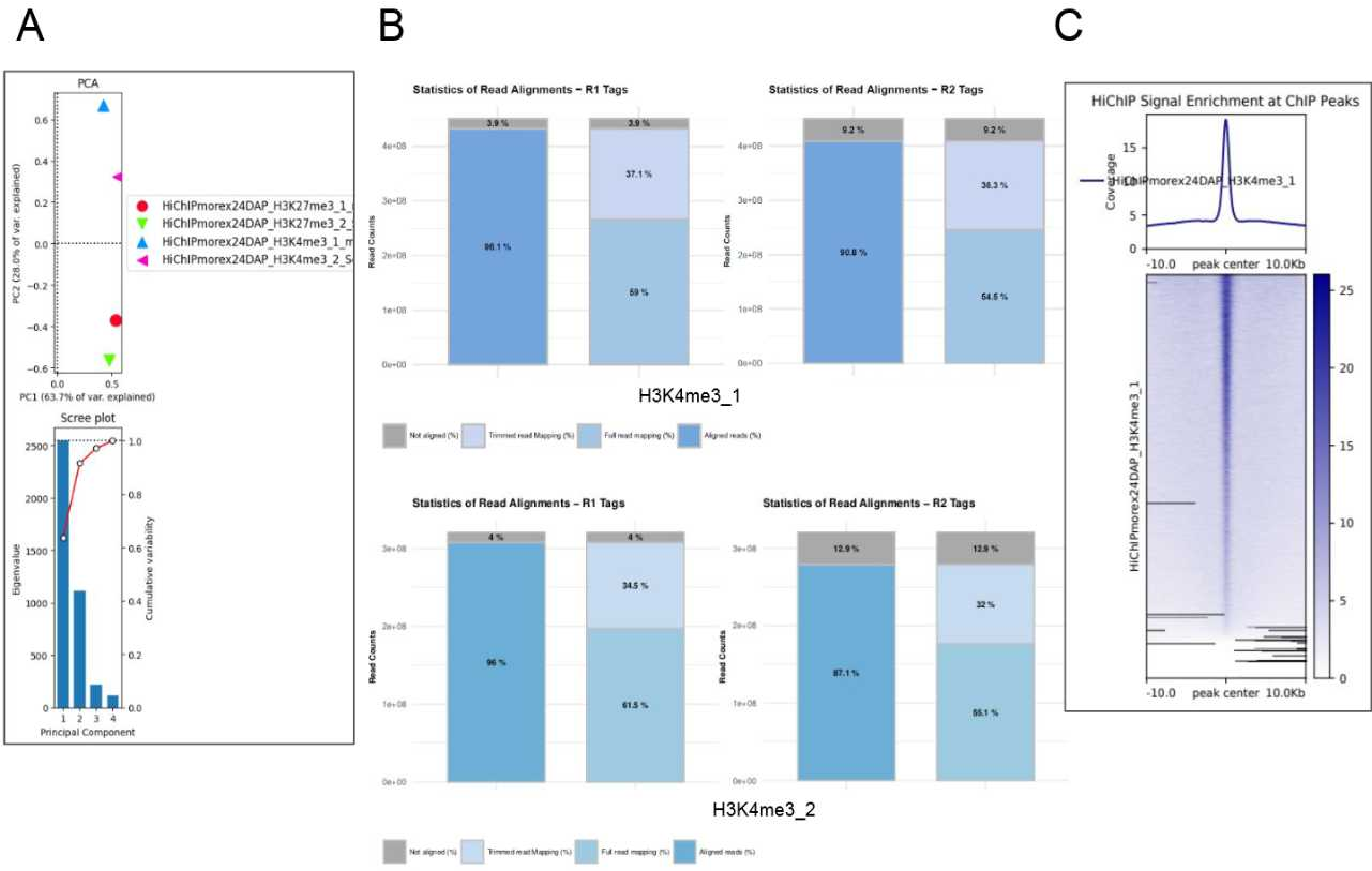


**Suppl. Fig. 6. Quality Control of the 24DAP H3K4me3 HiChIP sequencing datasets.** (A) HiChIP replicate correlation of mapped data by Principal Component Analysis (PCA). (B) Mapping statistics of the two 24DAP H3K4me3 HiChIP replicates. (C) HiChIP signal enrichment at the H3K4me3 ChIP peaks as called from previously performed native ChIP-seq experiment.

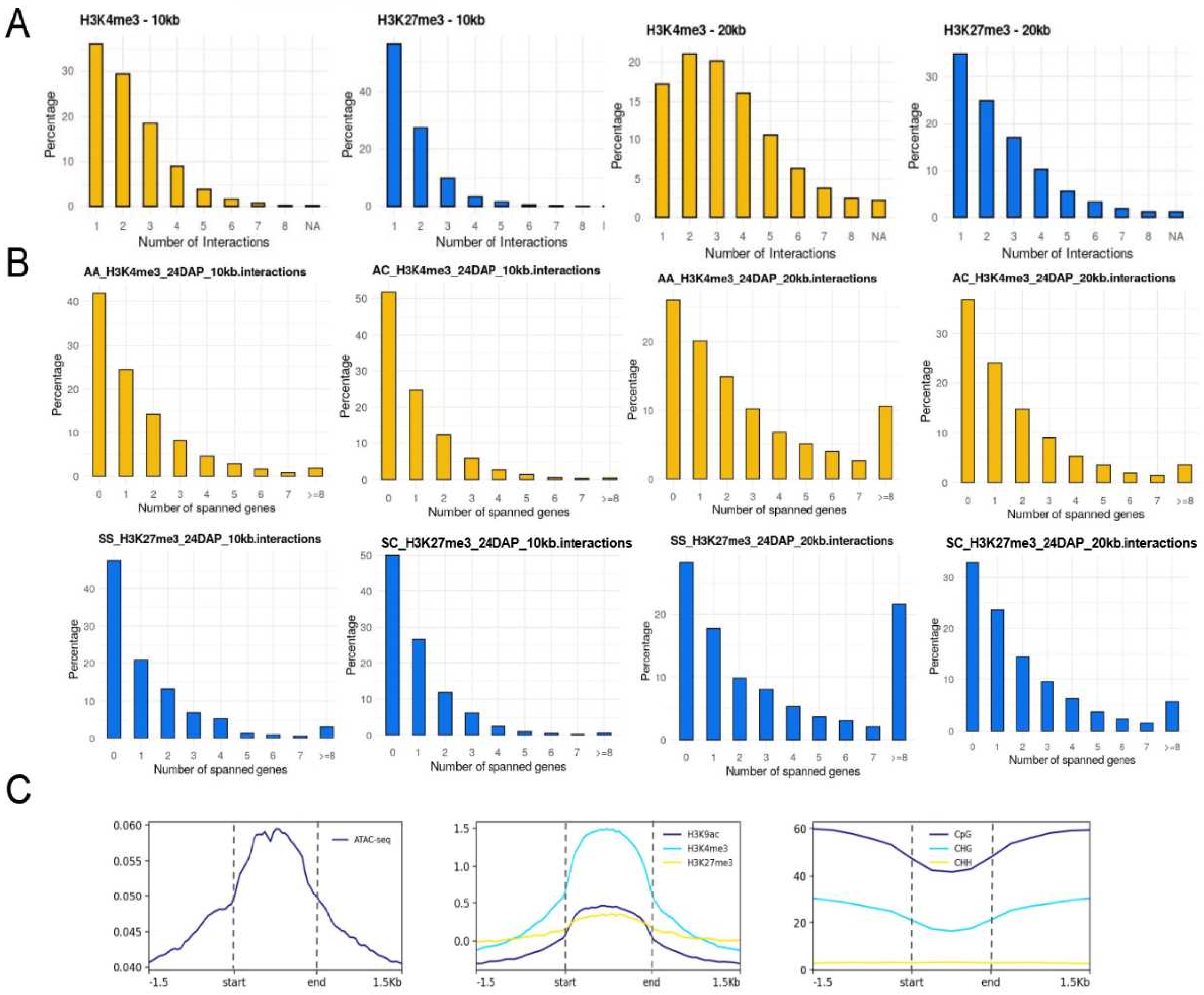


**Suppl. Fig. 7. Analysis of the interactome in the barley embryo.** (A) Numbers of H3K4me3 and H3K27me3 HiChIP interactions per promoter at lower resolutions (10- and 20-kb). (B) Numbers of genes spanned by H3K4me3 and H3K27me3 HiChIP interactions at lower resolutions (gene distances). (C) Enrichments of selected epigenetic features across H3K4me3 HiChIP anchors at 5-kb resolution indicates that they are transcriptionally dense regions.

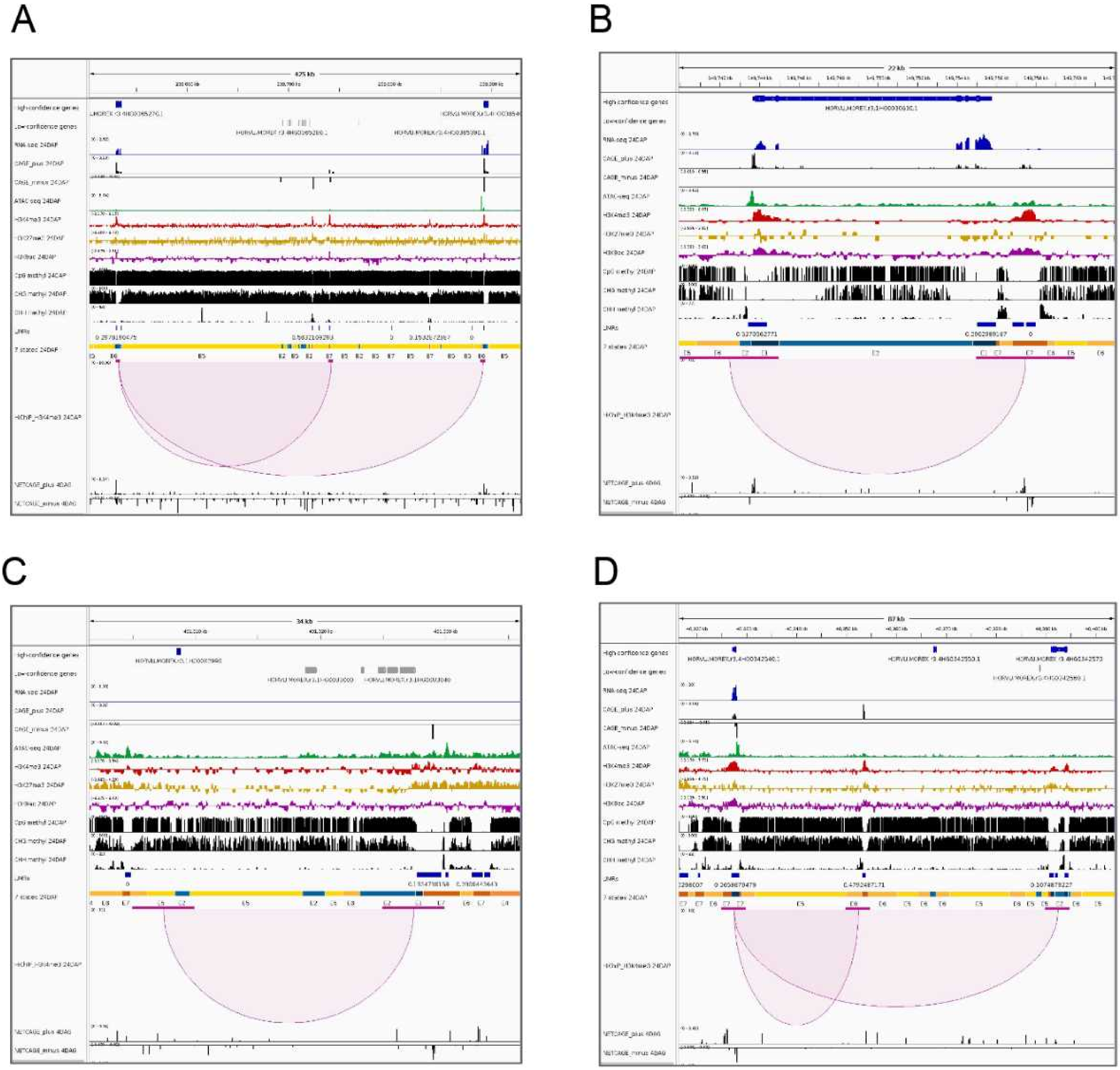


**Suppl. Fig. 8. Examples of distinct interaction classes together with epigenomic features** (A) Active promoter-active promoter, (B) active self-gene loop, (C) silent promoter-cCRE, (D) active promoter-cCRE and active promoter-silent gene.

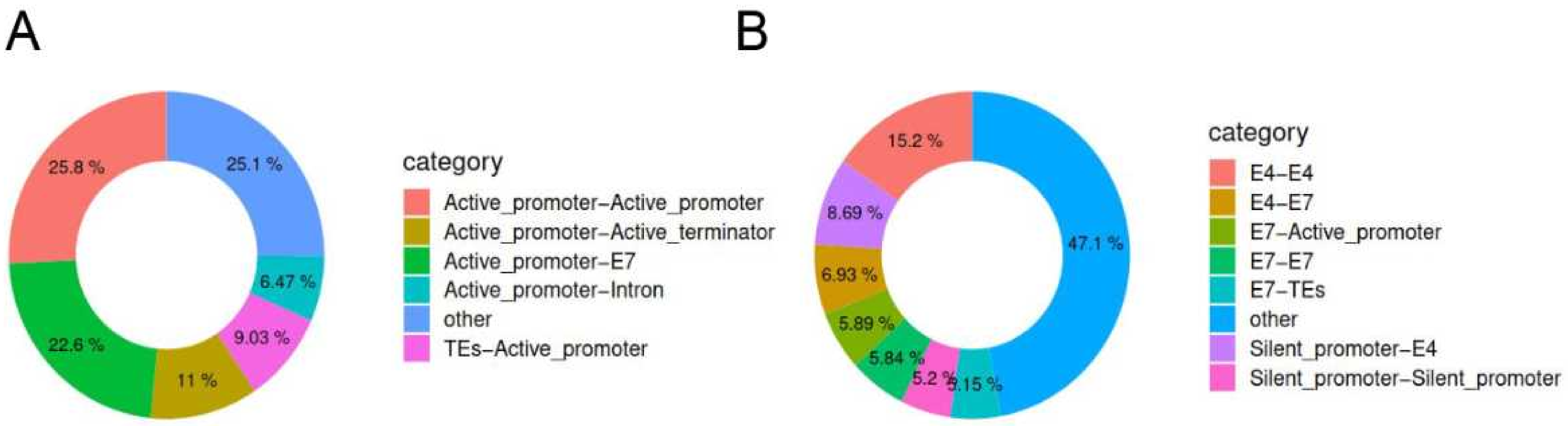


**Suppl. Fig. 9. Proportions of interaction classes identified by HiChIP at 5-kb resolution in 24DAP sample, focusing on high-confidence genes.** Annotation of all significant interactions associated with (A) activating (H3K4me3) histone mark and (B) repressive (H3K27me3) mark. The category 'other' includes all interaction classes with <5% frequency.

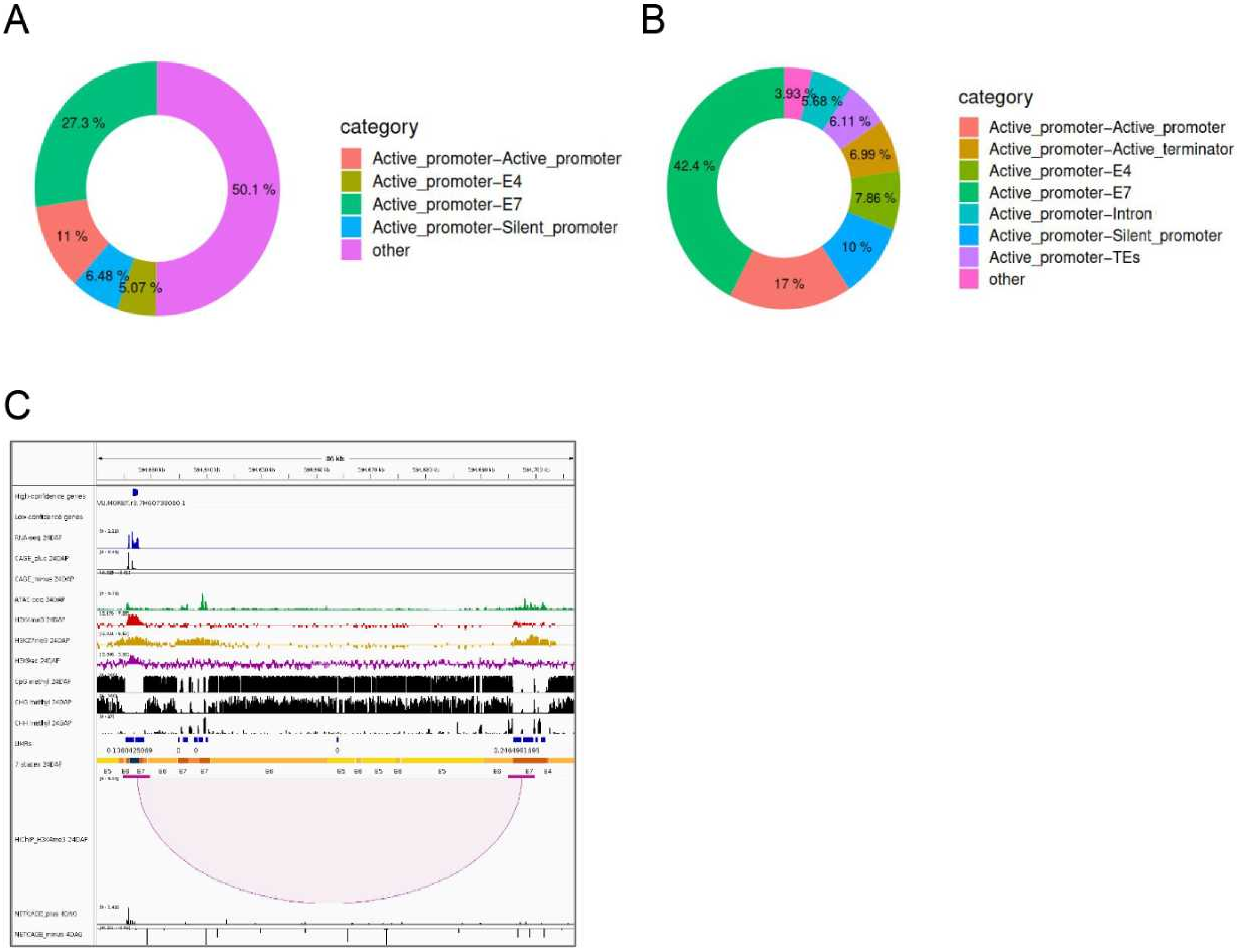


**Suppl. Fig. 10. Features of bivalent interactions.** Annotation of all bivalent interactions (A), and the same set from the active promoter-centric view (B). The category ‘other’ includes all interaction classes with <5% frequency. (C) An example of a bivalent interaction.
